# Neuroscience in Conversation: Tracking Scientific Change Through an Open Seminar Archive

**DOI:** 10.64898/2026.10.22.761073

**Authors:** Irene Bellin, Lena K. Tulner, Ahmad Beyh, Alfie Wearn, Stephanie J. Forkel

**Affiliations:** Donders Institute for Brain, Cognition and Behaviour, Radboud University, Nijmegen, The Netherlands; Institute for Interdisciplinary Studies, University of Amsterdam, The Netherlands; Department of Psychiatry, Brain Health Institute, Rutgers University, Piscataway, NJ, USA; Montreal Neurological Institute, Department of Neurology and Neurosurgery, McGill University, Montreal, QC H3A 2B4, Canada; Max Planck Institute for Psycholinguistics, Nijmegen, The Netherlands; Department of Psychology, School of Human and Community Development, University of the Witwatersrand, Johannesburg, South Africa

**Author notes:** These authors contributed equally to this work (Irene Bellin, Lena K. Tulner).

**Keywords:** Neuroimaging, science communication, open educational resources, YouTube, longitudinal corpus analysis, topic modelling, scientific discourse, open science

## Abstract

Scientific talks capture how researchers frame questions, debate ideas, and communicate emerging methods, providing a complementary record of scientific change to the published literature. We investigated whether an open neuroscience seminar archive could reveal shifting scientific attention and contain predictive information about subsequent discourse. We analysed 470 English-language transcripts from the Clinical Neuroanatomy Seminars YouTube channel, spanning 2010–2026 and approximately 2.6 million words, using longitudinal vocabulary analysis, unsupervised topic clustering, conceptual comparisons, and rolling temporal forecasting. The archive captured sustained discussion of language, white matter anatomy, connectivity, clinical neuroscience, and neurobiological mechanisms, alongside changes in their relative prominence. Network-related terminology became more prominent than localisation-related terminology, references to individual variability and computational approaches increased, and open science practices became more visible in scientific discussion. Beyond retrospective description, a regression model predicted keyword frequencies one and two years ahead with approximately 12% and 17% lower mean absolute error, respectively, than a persistence baseline. This modest gain is better explained by smoothing of year-to-year fluctuations than by genuine prediction: terms showing the strongest recent growth tended to fall back rather than continue rising. These findings suggest that spoken scientific discourse shows clear longitudinal change but offers limited evidence of information about subsequent shifts in attention. Interpretation remains grounded in the archive’s curated scope and changing content composition. By making nearly two decades of freely accessible neuroimaging education amenable to systematic analysis, this study connects open educational practice with the study of science itself. Open seminar archives can serve as reusable research resources, preserving the conversations through which scientific communities develop and revise their ideas.

## Introduction

Scientific communication preserves both research findings and the ways in which researchers explain, contextualise, and debate them. Digital platforms have expanded access to these exchanges, allowing scientific talks, methodological tutorials, and educational courses to reach audiences beyond the institutions and events in which they originated (Brossard, 2013; Wearn et al., 2025). The COVID-19 pandemic accelerated reliance on online educational resources: research in chemistry education documented a substantial increase in students’ use of instructional YouTube videos during this period (Breslyn & Green, 2022). These collections serve an immediate educational purpose while preserving scientific conversations that would otherwise leave little lasting record. As recordings accumulate, they also create an opportunity to investigate scientific discourse longitudinally: which questions attract attention, how explanatory frameworks are articulated, and how methods and research practices enter a community’s conversations (Blei & Lafferty, 2006).

Clinical and cognitive neuroscience draw on imaging methods with different histories and purposes, from CT and structural MRI for characterising anatomy and pathology to PET and functional MRI for investigating physiological and molecular processes. Continuing developments in acquisition, computational analysis, and multimodal integration rapidly expand their applications and reshape their interpretation (Boulant et al., 2024; Nozais et al., 2021; Tournier et al., 2011). Accessible expert instruction is therefore needed both to introduce new techniques and to explain the evolving capabilities and limitations of established approaches.

Free educational resources can reduce financial barriers to obtaining teaching materials, a concern highlighted in undergraduate neuroscience education (Hutchins & Harris, 2025). Open educational initiatives can remove subscription and membership barriers, although equitable participation also depends on language, connectivity, and accessible design (Levitis et al., 2021; Wearn et al., 2025). Discussions of neuroscience in Latin America and Africa further highlight the importance of addressing region-specific barriers and supporting locally grounded scientific expertise *(Besharati & Akinyemi, 2023; Silva et al., 2022; Francis et al., in press*). Access also depends on discoverability. Platforms such as World Wide Neuro (https://world-wide.org/) bring together neuroscience seminar listings and recordings from different institutions, complementing initiatives that produce and curate educational content. Together, these activities support access to scientific discourse beyond individual institutional networks.

The Clinical Neuroanatomy Seminars YouTube channel(@CNSeminars) exemplifies a sustained commitment to open neuroscience education. Originating in a departmental lecture series at King’s College London in the early 2000s, it launched on YouTube in 2016 to make specialised teaching and research in clinical neuroanatomy, neuroimaging, and clinical and translational neuroscience freely accessible (Guzmán Chacón et al., 2024). This transition preceded the COVID-19 pandemic by four years and subsequent efforts to open established neuroimaging archives such as the OHBM Time Machine initiative, which began making Annual Meeting recordings freely available in 2024 (Wearn et al., 2025). After six years of creating and digitising freely accessible educational content, the channel’s contribution was recognised with the 2022 Education in Neuroimaging Award from the Organization for Human Brain Mapping (OHBM) in Glasgow.

Over ten years, the channel expanded to encompass expert seminars, Master’s-level courses, methodological tutorials, Neuroccino journal clubs, and CNSdebates, alongside introductory videos and scientific documentary screenings. These formats combine instruction with opportunities to critically engage with findings, methods, and competing explanations. An earlier review documented the breadth of scientific discussion within the Neuroccino journal club, including debates about localisation, the ethical implications of artificial intelligence, and advances in neural decoding (Guzmán Chacón et al., 2024). Examples included a critique of localisationist explanations (Noble et al., 2024) and the reconstruction of music from human auditory cortex activity (Bellier et al., 2023). By August 2026, the channel contained 485 publicly available videos, including earlier recordings dating to 2010. The resulting archive therefore spans sixteen years of recorded content and preserves the changing interests of a community whose educational programme and scientific conversations developed over time.

For the ten-year anniversary of @CNSeminars in 2026, we investigate this archive as a measurable record of scientific attention in human cognitive neuroscience. Using 470 English-language transcripts, we characterise its thematic organisation and trace how topics, conceptual frameworks, methodological approaches, and research practices gain prominence, persist, or recede. We further test whether past vocabulary trajectories contain predictive information about subsequent frequencies through rolling temporal backtesting. Together, these analyses examine how an archive created for open education can also support systematic investigation of the development and communication of scientific ideas within an evolving research community.

## Methods

### Data and corpus definition

The primary data source for this study is the CNSeminars YouTube channel archive (https://www.youtube.com/@CNSeminars). As of the time of analysis (August 2026), the channel comprised 485 publicly available videos. Following eligibility screening (i.e., excluding videos without accessible transcripts and non-speech content such as silent recordings), 470 transcripts were retained for analysis. All retained content was in English.

An important distinction exists between recording year and upload year for a subset of videos. Although the channel launched on YouTube in 2016, several videos document talks and lectures delivered before then, with some content dating back to 2010. Analyses were indexed to the recording year where this information was available in the video metadata, and to the upload year otherwise, yielding a corpus spanning 2010 to 2026.

The channel hosts content of several distinct types, reflecting its evolution from a single lecture series into a broader scientific communication platform. The corpus includes CNSeminars talks (the channel’s founding and primary format), Neuroccino journal clubs, courses and educational series, community and open science content, and a residual category of other content. This typology was derived from video metadata and was used throughout the analyses to contextualise patterns in topic and terminology across content types.

### Transcript acquisition

Audio acquisition and transcription used a Python pipeline adapted from Jeffrey Emanuel’s *Bulk Transcribe YouTube Videos from Playlists* (https://github.com/Dicklesworthstone/bulk_transcribe_youtube_videos_from_playlist). The adapted implementation used pytubefix for audio retrieval and faster-whisper for local transcription with Whisper large-v3 (Radford et al., 2022). Transcription used beam search with a beam size of 10 and voice activity detection enabled to filter non-speech portions of the recordings. The pipeline selected CUDA GPU execution with half-precision computation when available, otherwise using CPU execution with automatic selection of the computation type. No language was explicitly specified in the transcription call. For each recording, recognised segments were concatenated into a plain-text transcript. Segment-level metadata were exported in CSV and JSON formats, including start and end timestamps, recognised text, and average log probabilities. These outputs retained temporal information linking the transcripts to the source recordings and model-derived information that could support subsequent quality assessment. The script also applied sentence segmentation to individual recognised segments using spaCy’s en_core_web_sm model. Transcript quality was assessed using the segment-level average log probability scores exported by the transcription pipeline. Transcripts with insufficient coverage, defined as fewer than 50 preprocessed tokens after cleaning, were excluded, as were recordings identified as containing predominantly non-English content. The latter were detected through automated token-level screening: a small number of transcripts (n = 6) contained substantial Welsh-language content arising from bilingual speakers, which could not be reliably handled within the English-language preprocessing pipeline and were excluded from all analyses.

Text preprocessing followed a multi-stage pipeline applied uniformly across all retained transcripts. Raw text was lowercased and cleaned of typographic artefacts including smart quotes and hyphenation variants. A compound-term joining step was applied prior to tokenisation, in which neuroscientifically meaningful multi-word expressions (for example, *white matter*, *corpus callosum*, *working memory*, and *default mode network*, etc.) were identified and concatenated with underscores to preserve them as single tokens throughout all downstream analyses. Tokenisation used a regular expression filter retaining only alphabetic tokens of four or more characters, removing numerals, punctuation, and isolated characters. Lemmatisation was applied using the NLTK WordNet lemmatiser.

A three-layer stopword scheme was applied iteratively and refined through manual inspection of the highest-frequency terms in the corpus. The base layer covered standard English function words. A seminar-specific layer removed high-frequency but non-discriminative terms characteristic of recorded academic talks, including platform logistics terms (e.g. *folder*, *joining*, *watching*), hedging and filler speech (e.g. *basically*, *actually*, *exciting*), and speaker names that appeared with sufficient frequency to distort term salience. A non-English layer removed tokens identified as Welsh or otherwise non-English, which appeared in a small number of transcripts containing mixed-language content despite the exclusions described above. Additionally, an automated consonant-cluster filter was applied to catch remaining non-English tokens not covered by the explicit stopword lists, discarding any token composed entirely of consonants after underscore removal. All stopword layers are defined explicitly in the analysis code and are fully reproducible.

### Longitudinal corpus analysis

To characterise how neuroscientific language has evolved across the decade, we applied a suite of corpus-level analyses to the temporally indexed transcript collection. Term frequencies were calculated for each year and normalised by the total corpus size for that year to account for substantial variation in the number of videos per year across the archive. Term frequency–inverse document frequency (TF-IDF) weighting was additionally applied to identify terms that are salient within a given year relative to their prevalence across the full corpus, providing a measure of terminological distinctiveness over time.

Vocabulary change was assessed by tracking the year of first appearance and last appearance of terms across the corpus, enabling identification of terms entering or leaving active scientific discourse. Temporal trajectories were modelled for a curated set of terms of theoretical interest, including established neuroimaging concepts, methodological terms, and emerging topics, with trajectory shape (growth, decline, plateau, resurgence) characterised descriptively and, where appropriate, by model fit to simple parametric forms. Terms exhibiting sustained growth or decline over consecutive years were classified as emerging or declining, respectively.

### Semantic embedding and topic clustering

To characterise the thematic structure of the corpus without imposing a fixed number of topics, each transcript was represented as a TF-IDF vector (scikit-learn’s TfidfVectorizer; Pedregosa et al., 2012), with a maximum vocabulary of 3,000 terms and a minimum document frequency of two transcripts. Vectors were normalised to unit Euclidean length before dimensionality reduction. Transcripts with fewer than 50 tokens after preprocessing (n = 4) were excluded, leaving 466 documents.

Dimensionality reduction was performed using Uniform Manifold Approximation and Projection (UMAP; McInnes et al., 2018), with cosine distance, 20 neighbours, and a minimum distance of 0.05, projecting each transcript vector into two dimensions. Clustering was performed in the reduced embedding space using HDBSCAN (Campello et al., 2013), a density-based algorithm that does not require a pre-specified number of clusters and labels documents not assigned to a cluster as noise. The minimum cluster size was 15, and the minimum number of samples was three.

Topics were labelled by extracting the highest-frequency preprocessed terms within each cluster. Topic prevalence by year was computed as the proportion of each year’s transcripts assigned to each cluster.

### Temporal topic dynamics and emergence topic prevalence over time

To examine how specific conceptual frameworks, methodological approaches, and scientific norms have evolved across the channel’s lifespan, we conducted a series of structured keyword analyses. Each analysis tracks the summed per-1,000-token frequency of predefined term sets across the full temporal range of the corpus, producing annual frequency trajectories that can be compared directly across term categories and time periods.

The first set of analyses targeted the relative prominence of contrasting conceptual and analytical approaches within human neuroscience. Five contrasts were defined a priori: localisation versus network and connectivity-based framing (Thiebaut De Schotten & Forkel, 2022); group-level inference versus individual differences and variability (Forkel et al., 2026); classical null-hypothesis testing versus Bayesian and machine learning approaches; task-based paradigms versus resting-state designs; and structural morphological description versus functional and dynamic characterisation. For each contrast, annual trajectories for both term sets were plotted, and crossover points (years in which one term set became more frequent than the other) were identified automatically. These contrasts were used to describe changes in vocabulary emphasis, without assuming that one approach superseded another.

The second set of analyses characterised the changing methodological and demographic composition of the corpus. Separate term sets were defined for neuroimaging modalities (i.e. fMRI, dMRI and tractography, EEG and MEG, fNIRS, PET, and non-invasive brain stimulation), study populations (healthy adults, clinical populations, developmental and paediatric samples, ageing, and sex/gender), and computational resources and tools (major open datasets, neuroimaging software, and deep learning frameworks). Annual frequency trajectories were examined per category to identify modalities gaining or losing prominence over the channel’s lifespan.

The third analysis examined the prominence of open science vocabulary within the channel’s scientific discourse. Term sets were defined for five dimensions: open science and open access; preregistration and preprints; reproducibility and replication; statistical reform; and FAIR data and sharing practices (Wilkinson et al., 2016). Annual trajectories were plotted, and mean frequency was compared between the early (2010–2017) and late (2018–2026) corpus periods to quantify the overall growth of open-science vocabulary. All term sets are defined explicitly in the analysis code and can be extended to track additional concepts.

### Emerging-topic forecasting and temporal validation

We asked whether the way a term’s prominence changed up to a given year helps to predict how it changes in the following one or two years. We call the last year of data available to the model the forecast origin *t*, and the years to be predicted *t*+1 and *t*+2. All transcripts from the same year were pooled into one document, giving 15 annual documents (2010–2017 and 2020–2026). No transcripts exist for 2018–2019, so 2017 and 2020 were treated as consecutive years. For each year we computed TF-IDF weights for the 800 most frequent terms occurring in at least two annual documents. Within each year, the weights were scaled so that the 800 values together had unit Euclidean length (L2 norm), which makes years of very different size comparable. A term’s value therefore describes its prominence relative to the other terms in the same year, not a raw count.

For each term and each forecast origin, we described the term’s history up to *t* with seven numbers: the slope of a straight line fitted over all years so far, the same slope over the last three years, the acceleration (the last-three-year slope minus the slope over the earlier years), the mean value over the last three years, the most recent value, the share of the term’s total value that falls in the last three years, and the proportion of years in which its value was above zero. A ridge regression (regularisation strength α = 1.0) was fitted across all terms at once, to learn how these seven numbers relate to a term’s value one year later (for *t*+1) or two years later (for *t*+2). To prevent information from the future from leaking into the model, it was trained only on examples in which both the descriptors and the value to be predicted came from years before the forecast origin, and it was then applied to the descriptors at the origin. Forecast origins began after four years of data.

Predictions were compared with the values actually observed (backtesting) and with two simple baselines: persistence (next year equals this year) and the three-year mean (the average of the last three years). We report the mean absolute error, which is the average size of the prediction error; the Spearman correlation between predicted and observed values; the rank skill, which is the Spearman correlation between predicted and observed change from the last observed value, computed across terms at each origin and averaged over origins; the number of origins at which the model’s error was smaller than that of persistence; and the directional hit rate, which is the proportion of terms for which the model correctly predicted whether the value would rise or fall (terms with no change were excluded), tested against 0.5 with a two-sided binomial test. Because the model predicted rises more often than they occurred, we used the rank skill, not the hit rate, as the main measure of directional accuracy.

Separately, we tested a score used to rank terms by recent growth. For a term with values v₁, …, v_t in the years up to the forecast origin *t*, the growth score was

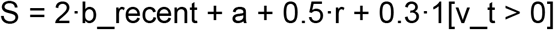

where b_recent is the slope of a straight line through the last three values, a = b_recent – b_early is the acceleration (b_early being the slope over the earlier values), r is the share of the term’s total value that falls in the last three years, and 1[v_t > 0] equals 1 if the term is present in the most recent year and 0 otherwise. The weights were set a priori to favour recent growth and were not tuned. At each forecast origin (2013–2025 for *t*+1 and 2013–2024 for *t*+2), we computed the score from the data up to that year and correlated it, across terms, with each term’s observed change in the following year or years. A positive correlation would mean that recent growth predicts further growth, and a negative one that recent risers fall back. Significance was assessed with a permutation test (2,000 shuffles of the observed changes across terms within each origin, using the mean correlation across origins as the test statistic), in both directions.

A ranking of terms by recent growth as of 2026 is shown as a descriptive result, together with the outcome of this test, and is not presented as a forecast. The framework is intentionally simple: with 15 annual observations the corpus is a short time series, and the aim is not precise forecasting but to establish whether any predictive signal exists and whether terms that are currently growing continue to grow.

### Sensitivity and robustness analyses

To test whether the clustering depended on the changing composition of the corpus, we re-ran the embedding and clustering pipeline on the CNSeminars videos only, on the corpus without Neuroccino, on the corpus without short transcripts (under 1,500 or 3,000 words), on the dense period 2020–2026, and on year-balanced subsamples (at most 20 videos per year). We also varied the clustering parameters: minimum cluster size, minimum samples, UMAP neighbours and minimum distance, vocabulary size, and the number of UMAP dimensions used for clustering. Because UMAP is stochastic, each scenario was repeated over ten random seeds and compared with the baseline run of the same seed; the seed-to-seed agreement of the baseline served as the reference. Agreement was quantified with the adjusted Rand index, the Jaccard overlap of matched clusters, and the correlation of their yearly prevalence trajectories (see Supplementary).

### Use of AI technologies

Transcripts of the video archive were generated with the automatic speech recognition model Whisper large-v3 (OpenAI; Radford et al., 2022), run locally through faster-whisper, as described under *Transcript acquisition*. The model’s output was screened for quality, language and length before analysis. Claude Sonnet 5.5 by Anthropic was additionally used to improve the language of the manuscript and assist with debugging analysis code. It was not used to generate data, results, figures or references. All AI-assisted output was reviewed, edited and verified by the authors. The authors are responsible for the accuracy, originality, attribution and integrity of the work, including all content produced or revised with AI assistance.

## Results

### Archive composition and Channel reach

The analytical corpus comprised 470 English-language transcripts spanning recordings from 2010 to 2026, with approximately 2.6 million words. Neuroccino journal clubs accounted for 240 transcripts, followed by CNSeminars talks (n=85), community and open science content (n=49), courses and educational series (n=46), and other content (n=50). Annual coverage was uneven, with a substantial increase in recordings from 2020 onwards (Fig. 1).

**Figure 1.**
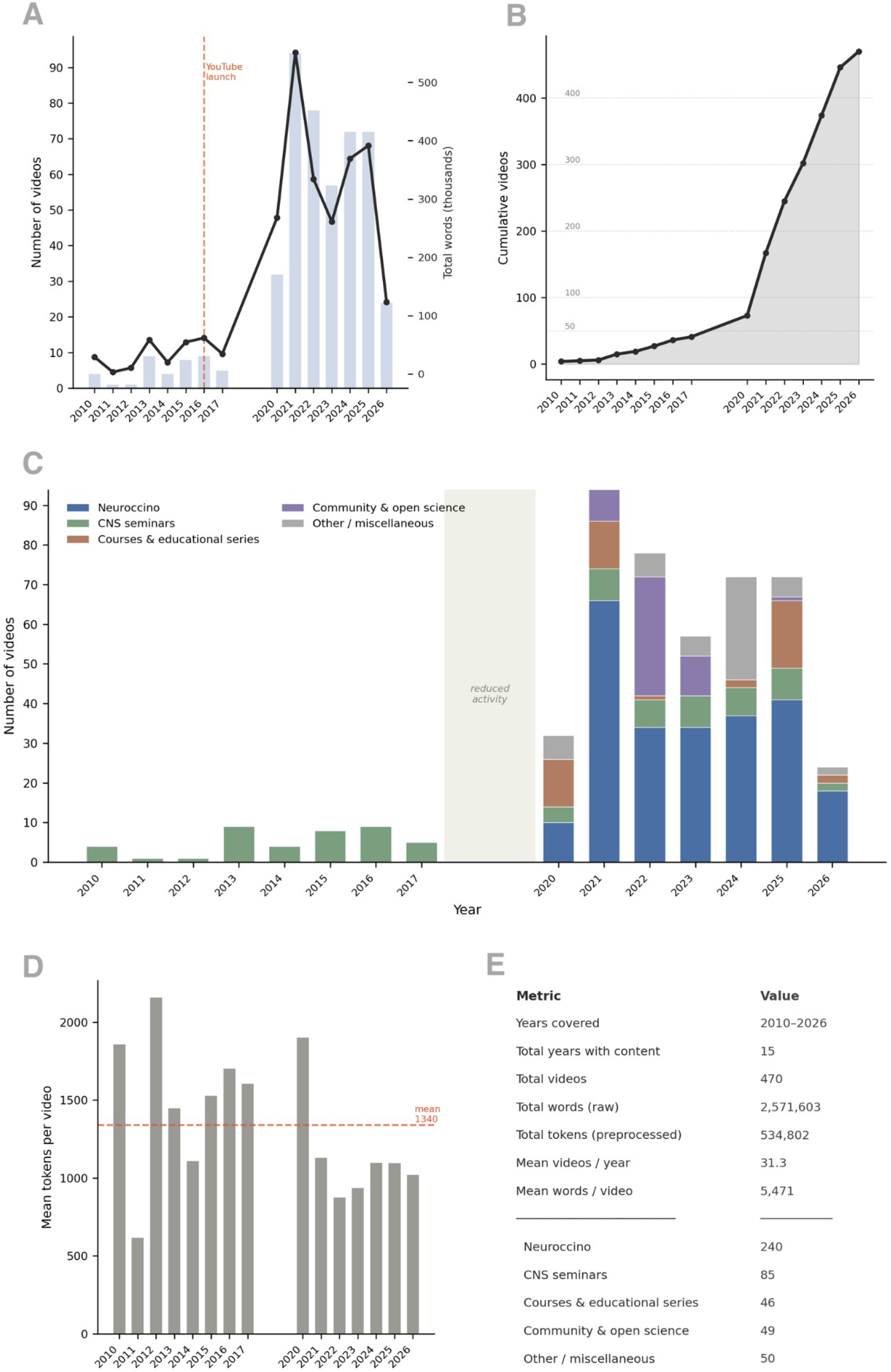
Overview of the CNSeminar video archive, 2010–2026. (A) Number of videos per year and total number of words per year. The dashed line marks the launch of the YouTube channel. (B) Cumulative number of videos, reaching 470 by 2026. (C) Number of videos per year by content type: Neuroccino, CNS seminars, courses and educational series, community and open science, and other or miscellaneous. The shaded band marks 2018–2019, for which no transcripts exist. (D) Mean number of tokens per video by year, after preprocessing. The dashed line shows the mean across years. (E) Summary of the archive. Before 2020 the archive consists only of CNS seminars; 2026 is a partial year.

**Figure 2.**
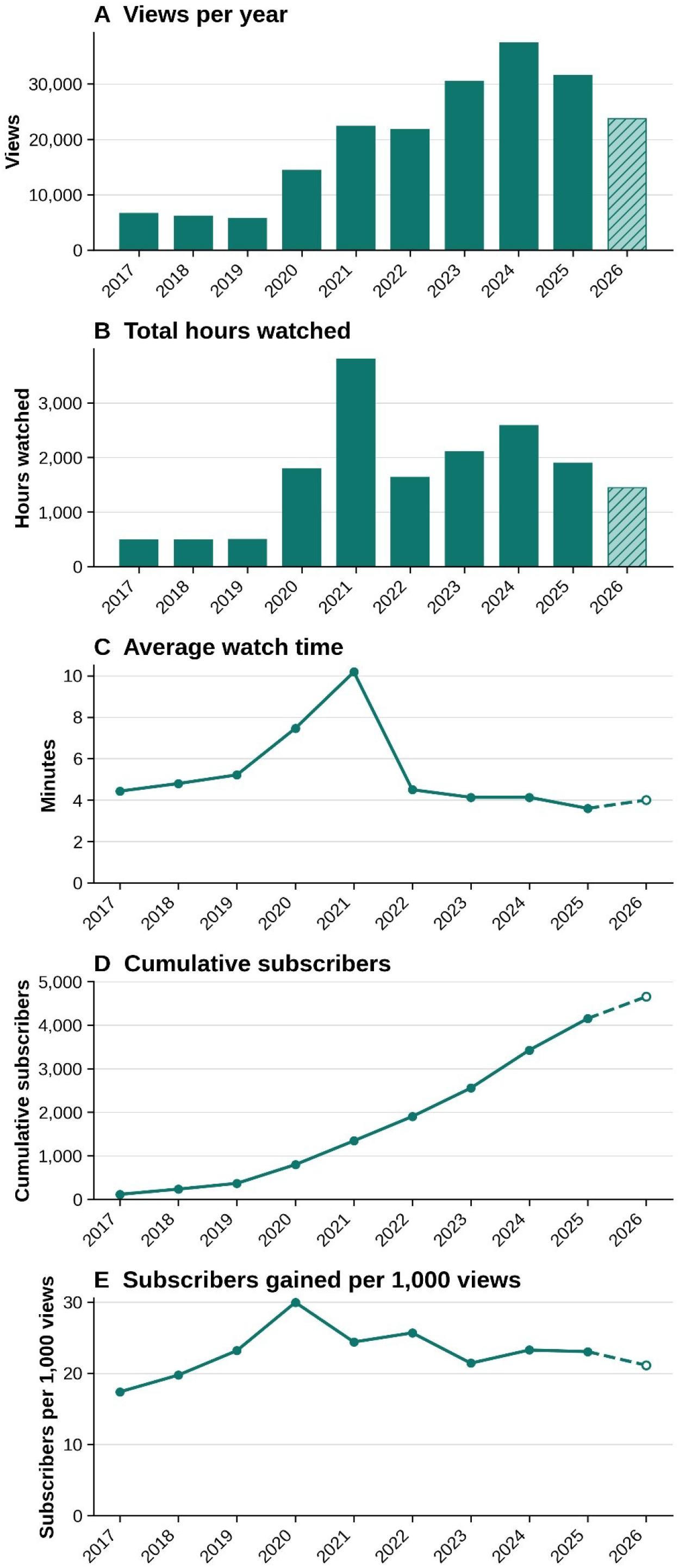
Channel-level engagement metrics of the CNSeminars YouTube channel, 2017–2026. (A) Annual views; (B) Total hours watched per year; (C) Average view duration (minutes); (D) Cumulative number of subscribers, calculated as the running total of annual subscriber gains; (E) Subscribers gained per 1,000 views in each year. Bars and lines show values per calendar year, as exported from YouTube Analytics. Hatched bars and dashed line segments indicate 2026, for which data are partial (data exported on 29 September 2026).

As of September 2026, the channel had approximately 4,690 subscribers. YouTube Analytics provided complementary information on the audience reached by the Clinical Neuroanatomy Seminars channel. Across the lifetime reporting period, viewers aged 25–34 and 35–44 accounted for 44.1% and 34.5% of views with available age data, respectively. The remaining age categories were 18–24 (11.2%), 45–54 (2.6%), and 65 years and older (7.6%). Within the platform’s gender categories, male and female viewers accounted for 61.1% and 38.9% of views, respectively. These distributions describe viewing activity for which demographic information was available, rather than the composition of all unique viewers. Recent audience measures indicated continued discovery of the channel alongside recurring engagement. As of 29 September 2026, the monthly audience was approximately 1,300 viewers, comprising 78.3% new, 16.8% casual, and 4.9% regular viewers. Among returning viewers during the preceding 28 days, 41% watched live content only, 37% watched recorded videos only, and 22% watched both formats.

Together, these measures document engagement with both synchronous and asynchronous neuroscience education. Between 25 February 2016 and 29 September 2026, the channel accumulated 203,249 views and 16,963.6 hours of watch time, with an average view duration of 5 minutes and 3 seconds. Geographic reporting documented viewing across Europe, Asia, Africa, the Americas, and Oceania. The United States contributed the largest reported country total (13,636 views), followed by Germany (2,979), France (1,965), and Italy (1,731). Viewing was also recorded in India, Brazil, and South Africa, as well as in Japan and Australia, indicating international reach beyond the channel’s institutional setting.

### Semantic landscape of the archive

Representing each transcript as a TF-IDF vector and projecting the vectors with UMAP revealed a structured thematic landscape (Fig. 3A). HDBSCAN resolved nine topic clusters containing 350 transcripts (75%), while the remaining 116 transcripts (25%) were not assigned to any cluster and lay mostly between clusters. The two largest clusters were T6 (neuron / cell / receptor; n = 89) and T7 (network / lesion / connectivity; n = 76). T6 formed an elongated, diffuse group at the centre of the map, which suggests a gradual continuum of cellular and systems-level neuroscience and not a sharply bounded topic. Language-related content was split into two neighbouring clusters in the upper left, T8 (language / speech / hemisphere; n = 37) and T9 (language / lesion / white matter; n = 26). Vocabulary on diffusion imaging and fibre anatomy formed two further clusters on the left, T5 (diffusion / fiber / axon; n = 30) and T4 (fiber / sulcus / cerebellum; n = 18). T1 (disease / gene / correlation; n = 28) and T3 (species / size / evolution; n = 17) were separate from the neuroanatomical core, and T2 (journal / code / open; n = 29) lay isolated at the bottom, consistent with its content on publishing, code and open science.

**Figure 3.**
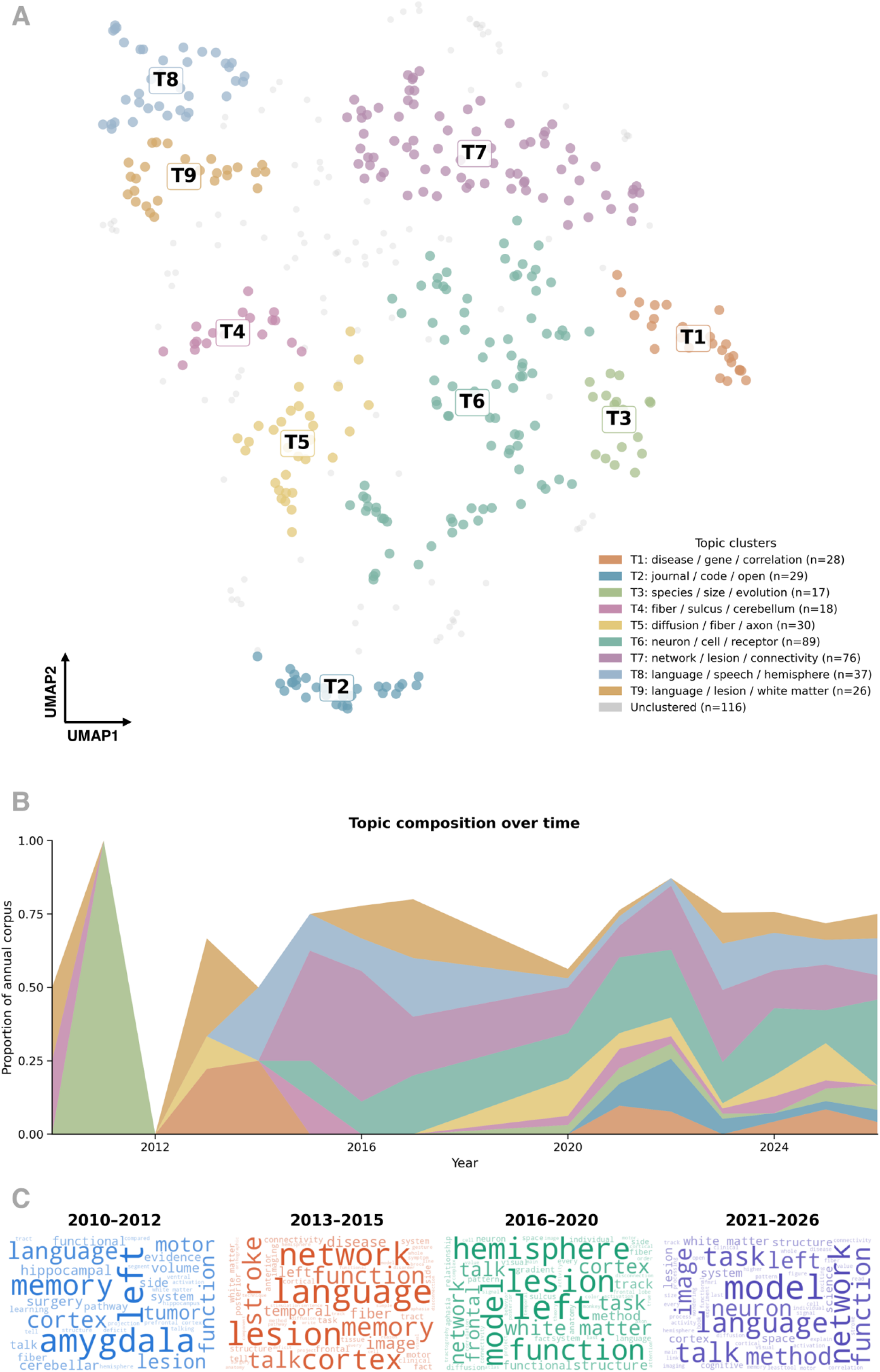
Thematic structure of the CNSeminar archive. (A) Two-dimensional UMAP projection of the TF-IDF vectors of all transcripts (n = 466); each point is one transcript. HDBSCAN identified nine topic clusters (T1–T9, colours), labelled with their three most frequent terms. 116 transcripts (25%) were not assigned to any cluster (light grey). (B) Composition of the annual corpus by topic, shown as the proportion of each year’s transcripts assigned to each cluster; the part of each year not covered by clusters is unclustered transcripts. Annual samples before 2020 contain between 1 and 9 transcripts (one each in 2011 and 2012), and the areas are interpolated across 2018–2019, for which no transcripts exist. (C) Word clouds of the most frequent terms in four eras (2010–2012, 2013–2015, 2016–2020, 2021–2026); word size is proportional to frequency.

The topic composition of the archive changed over time (Fig. 3B). Annual samples before 2020 were very small (1–9 transcripts per year), so the early composition, such as the single-topic year 2011 and the empty year 2012, reflects individual transcripts and should be read as descriptive only. From 2013, the language clusters (T8 and T9) and the network and lesion cluster (T7) made up a large share of the corpus until 2017, with T7 reaching roughly 0.4 of annual transcripts in 2015–2016. T6 (neuron / cell / receptor) was absent before 2015 and from 2021 onwards accounted for about a quarter of annual transcripts (0.2–0.3 per year), while T7 declined to below 0.1 in 2026. T2 (journal / code / open) first appeared in 2021 and peaked in 2022 at about 0.18. T5 (diffusion / fiber / axon) rose again towards 2025. Because unclustered transcripts are not shown, the stacked areas do not reach 1 in most years.

The vocabulary shows a corresponding shift (Fig. 3C). The earliest era (six transcripts) was characterised by regional and clinical-anatomical terms such as *amygdala*, *memory*, *hippocampal*, *cerebellar* and *tumor*. In 2013–2015 (21 transcripts), *language*, *network*, *lesion*, *stroke*, *fiber* and *tract* were prominent. In 2016–2020 (46 transcripts), *hemisphere*, *lesion*, *left*, *model* and *white matter* dominated, and in 2021–2026 (397 transcripts), *model*, *network*, *language*, *task* and *neuron* were most prominent, together with *method* and *image*. A core of terms (*language*, *function*, *left*, *cortex*, *lesion*) persisted across all four eras, whereas *model*, *task* and *method* moved from minor to leading terms, and *neuron* appeared prominently only in the most recent era.

### Vocabulary growth

The vocabulary of the archive grew steadily with the number of videos (Fig. 4A). The cumulative number of unique word types reached about 25,800 by 2026, and the number of new types added per year was closely linked to the size of each year’s corpus (Fig. 4B): from fewer than 1,000 types in 2011 and 2012 (one transcript each), through about 2,000–4,000 types a year in 2010–2017, to roughly 8,000–10,000 types a year from 2020 onwards. The type-token ratio fell from 0.67 in 2011 to about 0.09 in 2021 and then rose slightly to about 0.2 in 2026 (Fig. 4C). Because this ratio declines mechanically as texts get longer, the pattern mainly reflects the growth of the annual corpus and does not indicate a change in lexical diversity.

**Figure 4.**
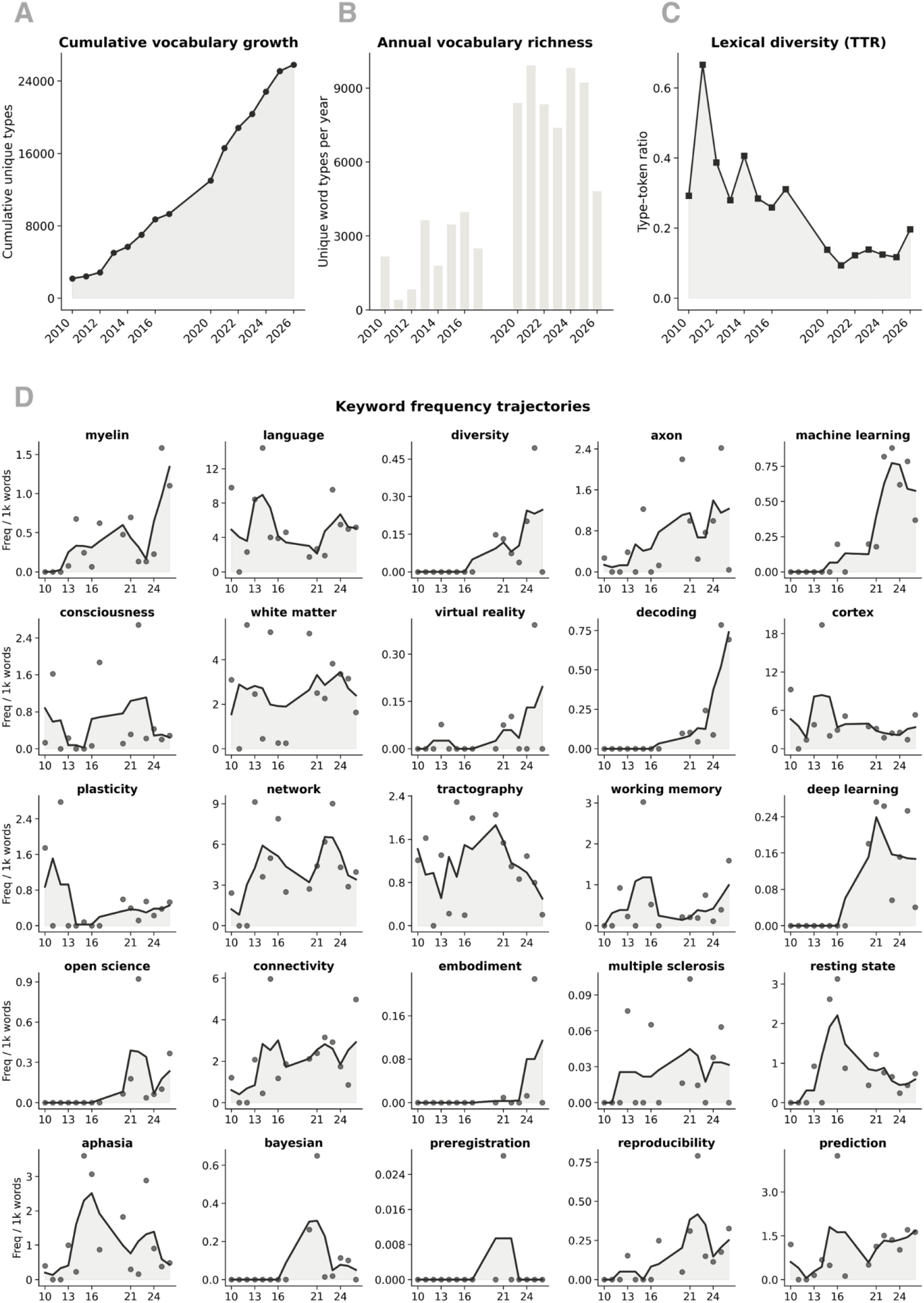
Vocabulary growth and keyword trajectories. (A) Cumulative number of unique word types in the archive, by year. (B) Number of unique word types in each year. (C) Type-token ratio by year (unique types divided by total tokens). (D) Frequency of 25 tracked terms per 1,000 preprocessed tokens. Dots are yearly values, and the line is a three-point moving average over consecutive observed years. No transcripts exist for 2018–2019, which were treated as consecutive steps on the time axis, and 2026 is a partial year. Years before 2020 contain between 1 and 9 transcripts.

Tracked terms showed several different trajectories (Fig. 4D). A first group emerged late and then rose. *Decoding* was near zero until about 2020 and rose sharply to about 0.7 per 1,000 tokens by 2025–2026; *myelin* and *axon* increased to about 1.3 and 1.2 per 1,000 tokens by 2026; and *diversity*, *embodiment* and *virtual reality* appeared only in the last few years, at low frequencies (below 0.25 per 1,000 tokens). *Machine learning* was absent before about 2016, rose from 2020, peaked in 2022–2023 at about 0.8 per 1,000 tokens, and then levelled off at about 0.6. *Deep learning* appeared in 2017 and peaked in 2021 (0.24). A second group rose and then fell. *Open science* and *reproducibility* peaked around 2021–2022 (0.4) and declined, *Bayesian* peaked in 2021 (0.3), and *preregistration* was practically absent throughout (below 0.03). *Aphasia*, *resting state* and *working memory* peaked around 2015–2016 and fell afterwards (*aphasia* from about 2.5 to 0.5, *resting state* from about 2.2 to 0.6), and *tractography* peaked in 2020 (1.8) and declined to about 0.5 by 2026. A third group remained present throughout without a clear trend, including *language*, *cortex*, *white matter*, *network* and *connectivity*; the first two were more frequent in the earliest years (*language* reached about 9 per 1,000 tokens in 2014), and *plasticity* was highest in 2010–2011. Because the early annual samples are small, the high early values of several terms rest on a handful of videos and should be interpreted with caution.

### Topics’ emergence, persistence and decline

To identify terms whose prominence changed systematically, we fitted a linear trend to the yearly TF-IDF weight of each of the 1,500 most frequent terms (Fig. 5). Twenty terms met the criteria for emerging terms (positive slope, R² ≥ 0.20; Fig. 5A). The steepest increase was for *neuron* (slope 0.0115 per year; R² = 0.69), well ahead of all other terms, which had slopes between about 0.004 and 0.007. They included other cellular and systems-level terms (*receptor*, *cell*, *mouse*, *monkey*), imaging and connectivity terms (*connectivity*, *functional connectivity*, *fmri*, *diffusion*, *gradient*, *atlas*), and general analytic terms (*prediction*, *variability*, *processing*, *cluster*, *condition*, *activation*, *cognitive*, *speech*). The trends with the best fit were those of *neuron*, *cluster* and *mouse* (R² = 0.69, 0.61 and 0.61), *variability* (0.55) and *cognitive* (0.51); for most other terms R² was between 0.20 and 0.40, indicating an upward but irregular trend. *Neurodesk*, the name of a software platform, also reached the list (slope 0.0065; R² = 0.20), which reflects a few sessions about that tool and not a field-wide trend.

**Figure 5.**
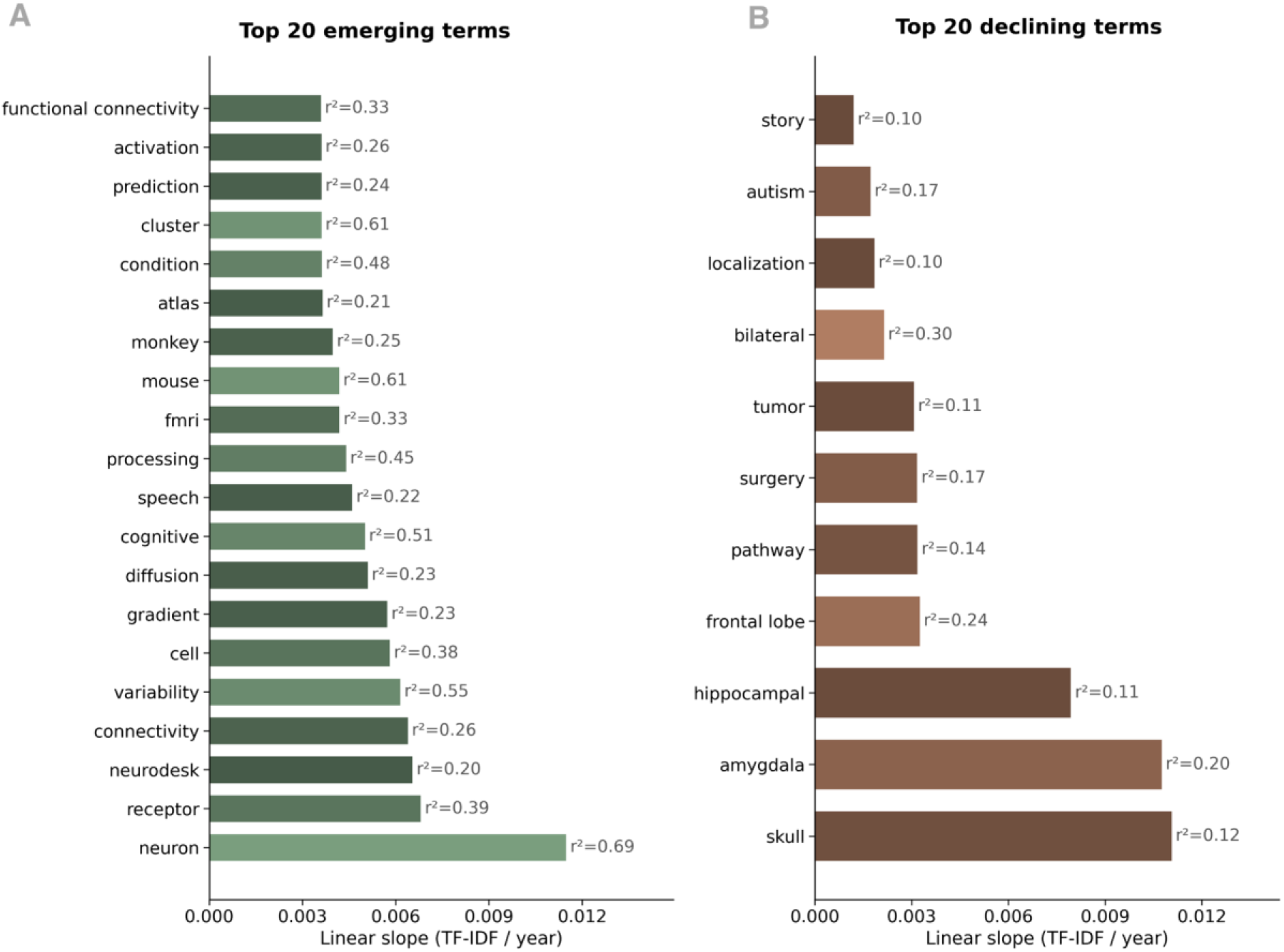
Terms with the strongest increase and decrease in prominence over time. TF-IDF weights of the 1,500 most frequent terms were computed on one document per year (L2-normalised within each year), and a straight line was fitted to each term’s trajectory across the 15 annual time steps (2018–2019 counted as one step). Only terms present in at least three years were considered. (A) The 20 emerging terms with the steepest positive slope (R² ≥ 0.20). (B) The 11 declining terms that met the criteria (negative slope, R² ≥ 0.10), shown as the size of the decrease. Bars show the linear slope, in TF-IDF units per annual step. The shade of each bar reflects the R² of the fit relative to the highest value in its panel (lighter = better fit), and R² is printed next to each bar.

Only eleven terms met the (more permissive) criteria for declining terms (negative slope, R² ≥ 0.10; Fig. 5B), and all of them fitted poorly (R² between 0.10 and 0.30). The steepest declines were for *skull* (slope −0.0111), *amygdala* (−0.0108) and *hippocampal* (−0.0079), followed by clinical and anatomical terms of smaller magnitude (*frontal lobe*, *pathway*, *surgery*, *tumor*, *bilateral*, *localization*), and by *autism* and *story*. The decreasing terms are therefore the clinical and regional vocabulary that characterised the earliest years of the archive (Fig. 4), whereas the emerging terms point to cellular neuroscience, connectivity and computational analysis, consistent with the growth of the neuron, cell, receptor and network clusters (Fig. 3).

Several caveats apply to these trends. Weights are relative to the other terms in each year, so a term can decrease because other terms increased. The early years contain very few transcripts (1–9 per year before 2020), so strong declines such as those of *skull* and *amygdala* rest on a small number of early videos, and their poor fit (R² ≤ 0.20) shows that they do not follow a steady downward trend. The increase of cellular terms also coincides with the growth of the Neuroccino series (about half of the corpus), which covers a broader range of topics than the earlier seminars.

### Conceptual shifts

To test whether the vocabulary of the archive reflects well-known conceptual shifts in neuroscience, we compared the frequencies of predefined terms for five pairs of paradigms (Fig. 6). The clearest shift was from localisation to networks. Terms of the networks paradigm (such as *network*, *connectivity*, *connectome* and *circuit*) overtook those of the localisation paradigm (such as *lesion*, *area*, *region* and *focal*) around 2011–2012 and stayed above them from then on. In the most recent years they were about three times as frequent (about 8 versus 2–3 per 1,000 tokens), while the localisation terms declined from a peak of about 8 per 1,000 tokens in 2016. Vocabulary on Bayesian methods and machine learning (*bayesian*, *machine learning*, *deep learning*, *prediction*, *decoding*, *classifier*) was also consistently more frequent than that of classical statistics (*anova*, *regression*, *t-test*, *p-value*), and it rose from about 1–2 per 1,000 tokens in the 2010s to about 3.8 in 2026, whereas classical terms stayed below about 0.6 throughout. In this case the two curves were never close, so the pattern describes the growth of one vocabulary and not the replacement of the other.

**Figure 6.**
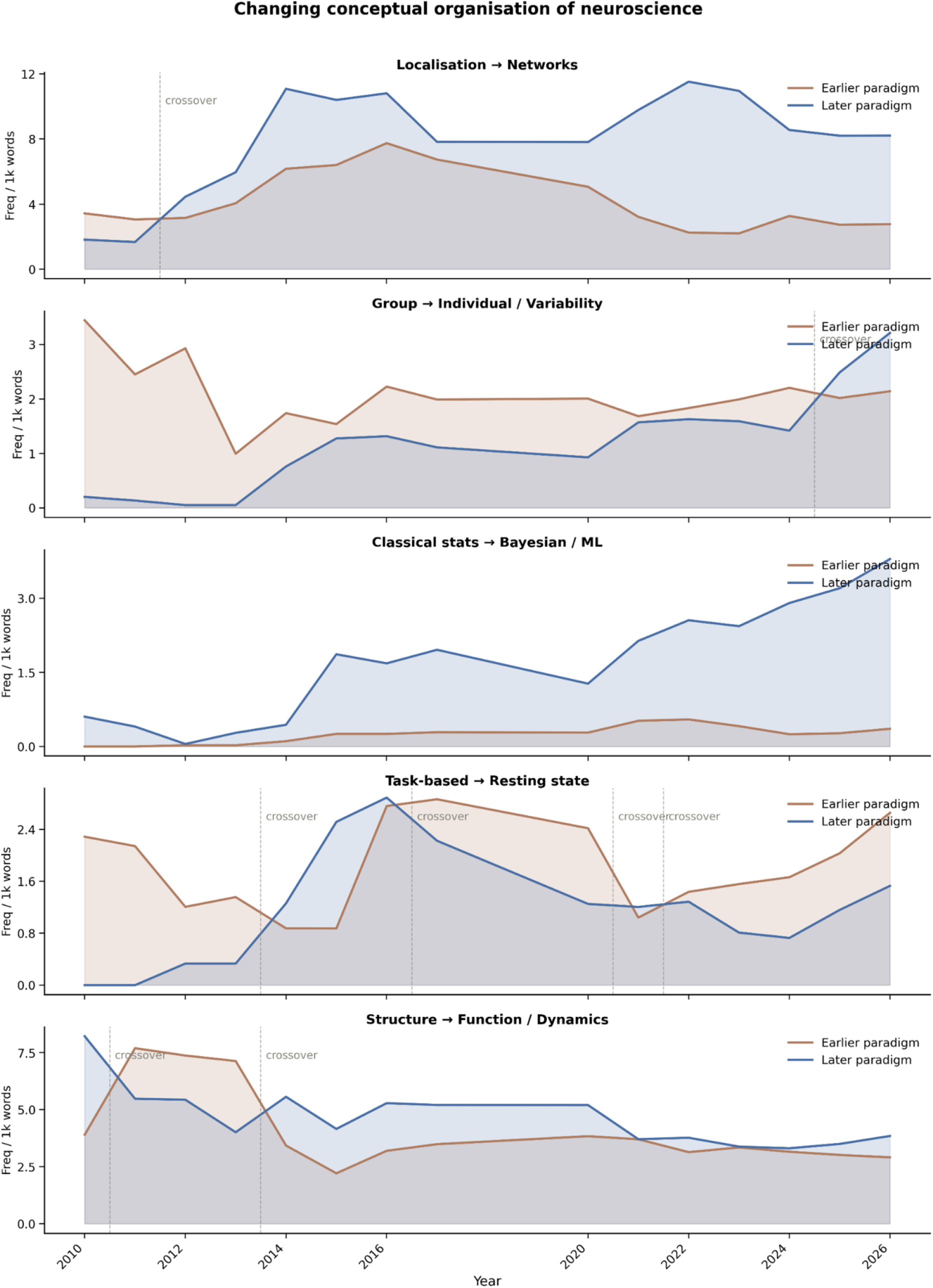
Conceptual shifts in the vocabulary of the archive. For each of five pairs of research paradigms, the lines show how often terms of the earlier paradigm (brown) and of the later paradigm (blue) occur in each year, as frequency per 1,000 preprocessed tokens, summed over all terms in the group and smoothed with a three-point moving average over consecutive observed years. The pairs are: localisation → networks, group → individual and variability, classical statistics → Bayesian and machine learning, task-based → resting state, and structure → function and dynamics. Dashed vertical lines mark years in which the two curves cross. The term lists were defined a priori. No transcripts exist for 2018–2019, and years before 2020 contain between 1 and 9 transcripts.

The shift from group to individual terms came late. Group-level vocabulary (*group*, *average*, *mean*, *population*) dominated until the last few years, and terms for individual differences and variability (*individual*, *variability*, *heterogeneity*, *precision*) rose steadily from the mid-2010s and overtook them only in 2025–2026 (about 2.5– 3.2 versus about 2.0–2.1 per 1,000 tokens). The shift from task-based to resting-state vocabulary did not show a lasting replacement. Resting-state terms (*resting state*, *default mode network*, *intrinsic*) briefly exceeded task terms around 2014–2016, but the two curves crossed several times, and task vocabulary was again more frequent in recent years (about 2.6 versus 1.5 per 1,000 tokens in 2026). Structure and function terms converged: function and dynamics vocabulary (*dynamics*, *temporal*, *oscillation*, *functional*, *activity*, *signal*) was higher than structural vocabulary (*anatomy*, *morphology*, *volume*, *thickness*) in most years, but the difference narrowed from about 2021 and both groups were close to 3–4 per 1,000 tokens in the latest years. Overall, the archive documents the rise of network and data-driven approaches, whereas other transitions are partial or still in progress.

### Changing methods, populations and resources over time

Vocabulary on imaging modalities was dominated by diffusion imaging and tractography throughout most of the archive (about 4–7 per 1,000 tokens in most years; Fig. 7A). MRI and fMRI vocabulary rose from about 3.4 per 1,000 tokens in 2017 to about 6.4 in 2026 and exceeded diffusion terms in the latest year (about 6.4 versus 4.9). The other modalities were less frequent. Vocabulary on TMS and tDCS peaked around 2016 (about 2.8) and fell to about 1 per 1,000 tokens in the 2020s. EEG and MEG terms increased in 2016 to about 2.4 and remained at that level until 2020 before falling below 1 per 1,000 tokens. PET terms stayed low (about 0.2–1.3) with a slight rise in the latest years, and fNIRS terms were practically absent in all years.

**Figure 7.**
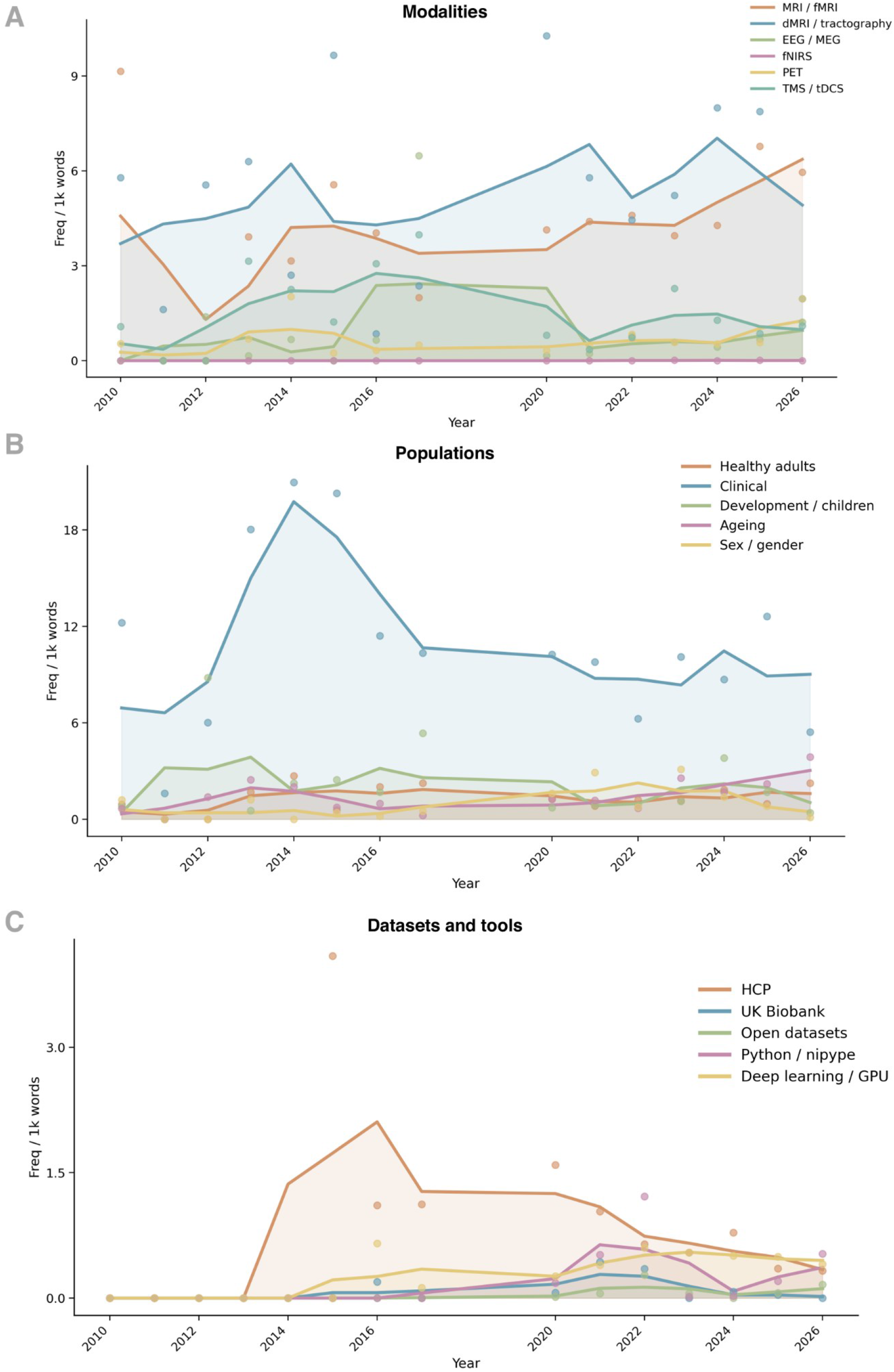
Methods, study populations and resources in the vocabulary of the archive. Each line shows the frequency per 1,000 preprocessed tokens of a predefined keyword set in each year; dots are the yearly values and lines are a three-point moving average over consecutive observed years. (A) Imaging and stimulation modalities. (B) Study populations. (C) Datasets and software tools. The frequency of a category is the sum over all its terms, so categories with more or more frequent terms have higher values and should be compared across time rather than with each other. No transcripts exist for 2018–2019, 2026 is a partial year, and years before 2020 contain between 1 and 9 transcripts.

Clinical vocabulary was by far the most frequent population category (Fig. 7B). It rose from about 7 per 1,000 tokens in 2010 to a peak of about 20 in 2014, declined to about 10 by 2017–2020, and remained at about 8–10 until 2026. The other categories stayed below about 4 per 1,000 tokens. Terms for development and children were most frequent in 2012–2016 (about 3–4) and then declined. Ageing terms increased gradually to about 3 per 1,000 tokens in 2026. Terms for sex and gender were highest around 2021–2022 (about 2.3) and decreased afterwards, and terms for healthy adults remained around 1–2 throughout.

Resource-related vocabulary was infrequent and changed over time (Fig. 7C). Terms for the Human Connectome Project appeared in 2014 and peaked in 2016 (about 2.1 per 1,000 tokens), then declined to about 0.3 in 2026. Terms for Python and neuroimaging software (such as *python*, *nipype* and *fsl*) appeared in 2017, peaked in 2021 (about 0.6), fell in 2024 and recovered slightly in 2026. Deep learning and GPU terms rose from 2015 to about 0.5 per 1,000 tokens in the 2020s. The UK Biobank and open datasets remained rare (at most about 0.3 per 1,000 tokens), so they are not informative as trends. Overall, the archive shows a stable emphasis on structural connectivity and clinical research, a recent increase in fMRI and ageing vocabulary, and a growing but small presence of computational tools.

### Open science era

Vocabulary related to open and reproducible science was almost absent in the early archive and increased markedly in the 2020s (Fig. 8). The mean annual frequency of the pooled open-science terms was about 0.23 per 1,000 tokens in 2010– 2017 and about 1.89 in the later period, roughly eight times higher (Fig. 8B). The groups of terms did not rise together (Fig. 8A). Vocabulary on reproducibility appeared first, from 2012, and rose from 2015 to about 0.5 per 1,000 tokens by 2017. Terms on statistical reform followed, increasing from 2014 to about 0.6 in 2021. In the 2020s, terms for open science and for FAIR data sharing grew most steeply. Open-science terms rose from about 0.2 in 2020 to a plateau of about 0.6 in 2021–2022, and FAIR and sharing terms rose to about 0.3 in 2021–2022. Reproducibility terms reached their highest value in 2022 (about 0.9 per 1,000 tokens). After 2022, open-science and reproducibility vocabulary declined, with a trough in 2024 (open science about 0.15; reproducibility about 0.4), and in 2025–2026 open-science and FAIR terms increased again (to about 0.3 and 0.45), whereas statistics-reform terms fell to about 0.25 in 2026. The group labelled preregistration remained the least frequent throughout (at most about 0.2 per 1,000 tokens). Overall, open-science vocabulary has become a regular, if small, part of the seminar discussions, but its use is uneven from year to year.

**Figure 8.**
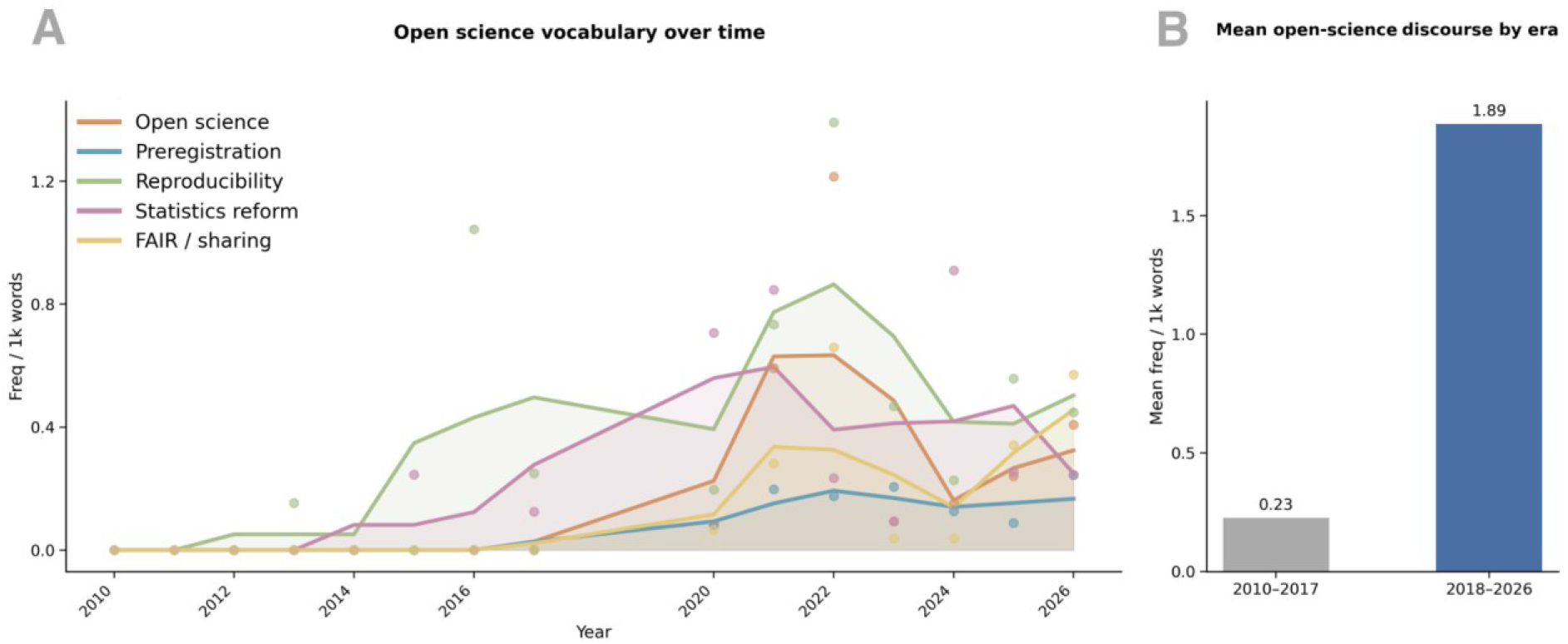
Open-science vocabulary increased over time. (A) Frequency per 1,000 preprocessed tokens of five groups of open-science terms in each year (open science, preregistration and preprints, reproducibility, statistics reform, and FAIR and data sharing). Dots are yearly values and lines are a three-point moving average over consecutive observed years. (B) Mean annual frequency of all open-science terms pooled, in the earlier period (2010–2017; 41 transcripts) and the later period (2020– 2026; 429 transcripts; no transcripts exist for 2018–2019). Each year contributes equally to the mean.

### Emerging topics prediction

In a rolling-origin backtest, the ridge model predicted term values better than the persistence baseline (Fig. 9). The mean absolute error was 0.0139 for the model and 0.0158 for the baseline at one year ahead (12% lower), and 0.0140 versus 0.0168 at two years ahead (17% lower; Fig. 9A). The model had the smaller error in 9 of 11 forecast origins at t+1 and in all 9 at t+2, and the advantage was largest for origins in the mid-2010s (Fig. 9B). Predicted and observed values were positively correlated (Spearman ρ = 0.45 at t+1 and 0.39 at t+2). However, the model’s predictions were compressed towards typical values (mostly between 0.01 and 0.07), and terms with high observed values were underpredicted (Fig. 9A). Part of the improvement over persistence therefore reflects shrinkage of extreme values towards the mean and not detection of trends. In line with this, a simple average of the last three years performed slightly better than the model (mean absolute error 0.0137 and 0.0138, against 0.0139 and 0.0140 for the model), and the directional hit rate (0.66 and 0.70; both p < 0.001 against chance) was inflated because the model predicted increases for 68% (t+1) and 70% (t+2) of terms that changed, while only 51% and 54% of these terms actually increased. The rank skill, the mean within-origin correlation between predicted and observed change, was 0.52 at t+1 and 0.54 at t+2. We therefore consider the model to have modest skill, mainly in smoothing noisy annual values and moderating extreme ones, and not in anticipating sustained growth or decline.

**Figure 9.**
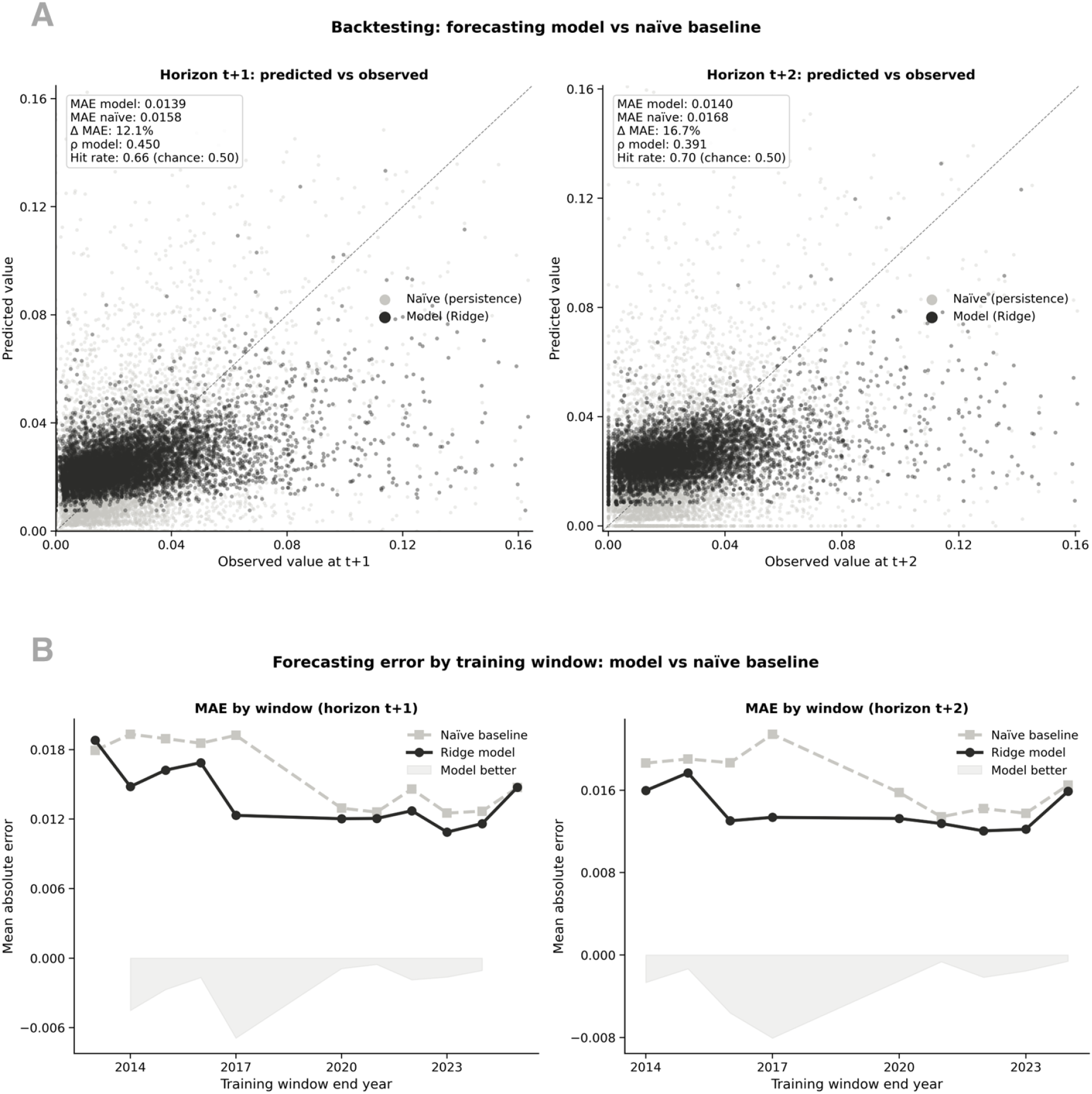
Backtest of the forecasting model. (A) Predicted versus observed term values at one year (t+1, left) and two years (t+2, right) ahead, pooled over all rolling forecast origins and 800 terms. Dark points are predictions of the ridge regression, light points are the persistence baseline (next year equals this year), and the dashed line marks perfect prediction. Values are year-level TF-IDF weights (L2-normalised within each year). The insets give the mean absolute error (MAE) of the model and of the baseline, the relative reduction in MAE, the Spearman correlation between predicted and observed values, and the directional hit rate. (B) MAE of the model (dark) and of the persistence baseline (light) for each forecast origin, plotted by the last year of the training window. The shaded area below zero shows the difference between the two (model minus baseline) where the model has the smaller error. The model was trained only on information available before each origin.

We ranked terms by recent growth (Fig. 10A). The highest-ranking terms as of 2026 were *hierarchy*, *receptor*, *neuron*, *balance* and *prefrontal cortex*, followed by *variability*, *monkey*, *aging*, *myelin*, *Alzheimer* and *fMRI*. To test whether such a ranking anticipates continued growth, we applied the same score at every earlier forecast origin and correlated it with the term’s subsequent change (Fig. 10B). The correlation was negative at every origin and both horizons (all 21 of 21). At one year ahead the correlations ranged from −0.48 to −0.21 (mean ρ = −0.34), and at two years ahead from −0.52 to −0.15 (mean ρ = −0.34); a permutation test indicated that this was very unlikely to arise by chance (p < 0.001). The terms that had grown most therefore tended to fall back in the following years, a pattern of regression to the mean in which a rise in a term’s prominence reflects the programme of a given year and not a sustained trend. For this reason, the ranking in Fig. 10A describes recent growth and should not be read as a forecast of emerging topics.

**Figure 10.**
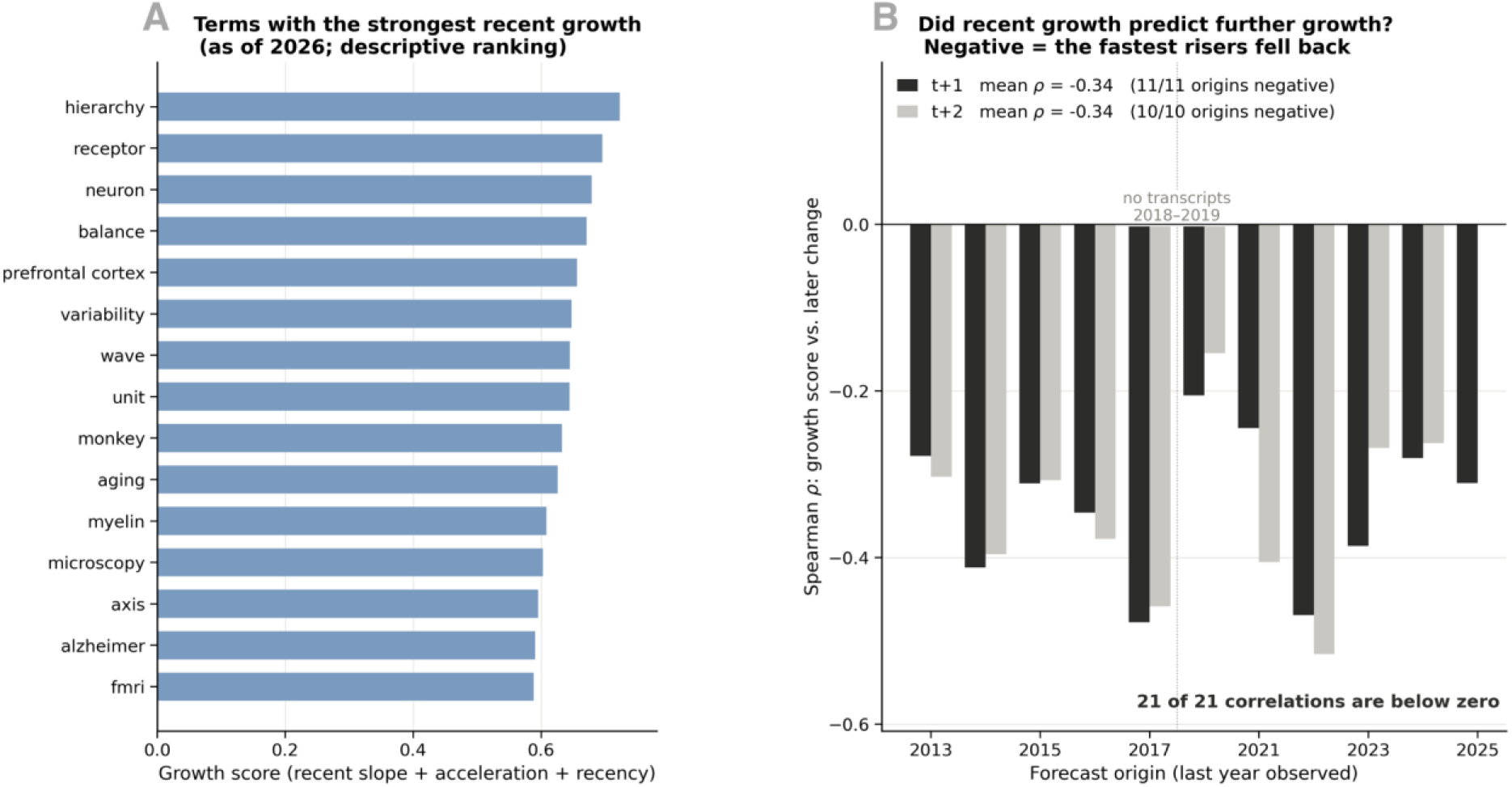
Terms with strong recent growth did not keep growing. (A) The 15 terms with the highest growth score as of 2026 (growth score = 2 × recent slope + acceleration + 0.5 × share of value in the last three years + 0.3 if present in the latest year). The ranking is descriptive. (B) Spearman correlation between the growth score and each term’s change one year (dark) and two years (light) later, across the 800 tracked terms, for every forecast origin from 2013 to 2025 (the last year observed). Negative values mean that the fastest-rising terms declined afterwards. All 21 correlations were negative (mean ρ = −0.34 at both horizons). The dotted line marks 2018–2019, for which no transcripts exist.

### Neuroccino journal selection bias

The papers discussed at Neuroccino sessions between June 2023 and June 2026 (n = 106) came from 45 sources, but they were concentrated in a few publishers (Fig. 11). Nature Portfolio Group journals accounted for 37 papers (35%), more than three times as many as any other group. The remaining papers were published on preprint servers (10; 9%), Elsevier journals (8; 8%), Oxford University Press (8; 8%), Cell Press (7; 7%), PLOS (6; 6%), PNAS (5; 5%) and Science / AAAS (5; 5%), with the other 10 groups contributing 1–3 papers each (Fig. 11A). At the level of single sources, *Nature Neuroscience* was the most frequent (10 papers), followed by bioRxiv and medRxiv (9), *Nature* and *Nature Communications* (7 each), *Brain* (6) and PNAS (5); the other 39 sources contributed three or fewer papers each (Fig. 11B). Thirty-one of the 92 papers in published journals (34%) appeared in a small set of high-profile journals (*Nature*, *Science*, *Cell*, *Neuron*, *Nature Neuroscience*, *Nature Reviews Neuroscience*, *Nature Human Behaviour* and *Nature Methods*). This composition indicates that the Neuroccino readings were selected mainly from prominent journals, in particular from the Nature family, and it means that the Neuroccino part of the corpus reflects the output and interests of these outlets and not the neuroscience literature as a whole.

**Figure 11.**
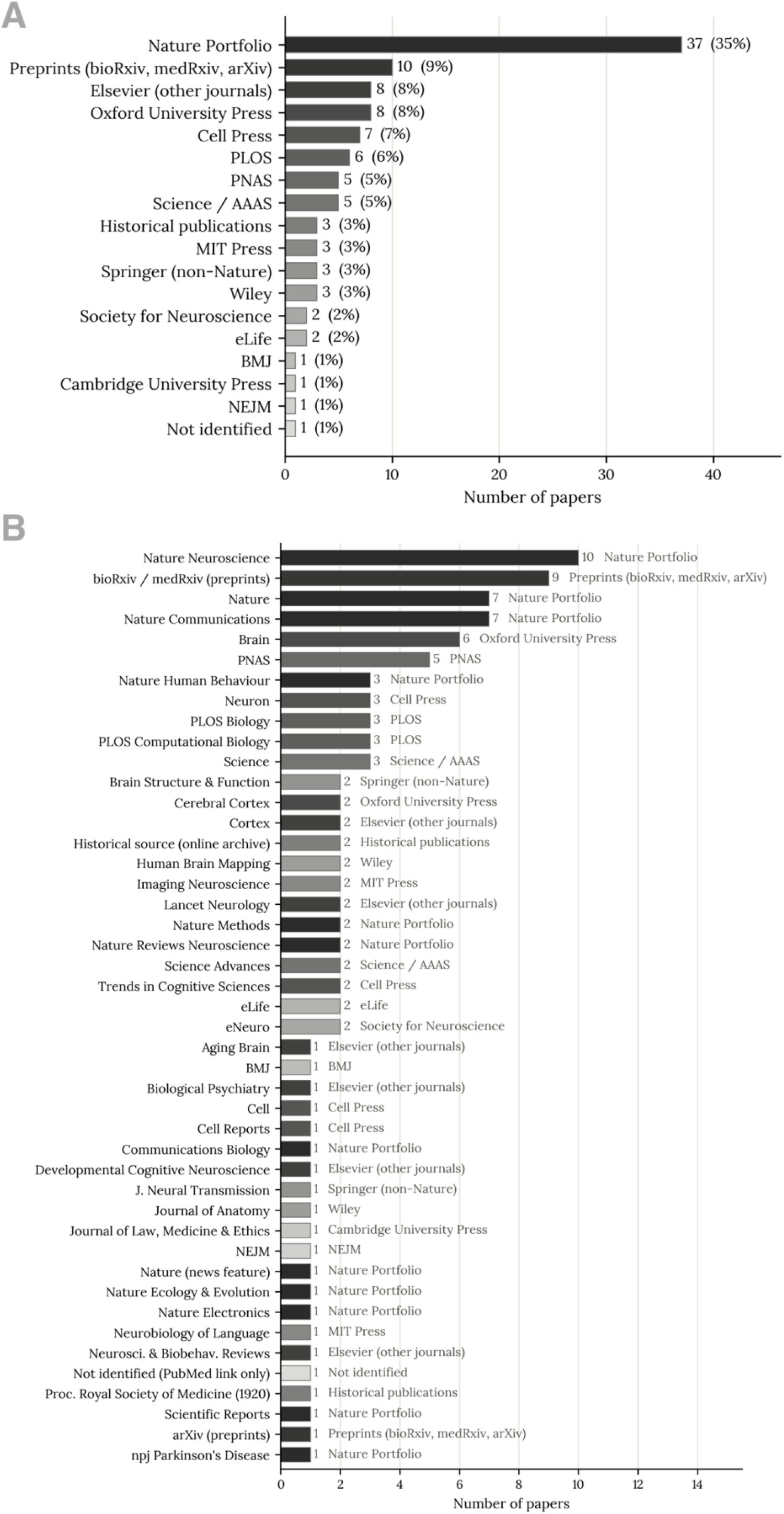
Journals and publishers of the papers discussed at Neuroccino sessions. (A) Number of papers by publisher or journal group (18 groups; percentages of n = 106). (B) Number of papers for each of the 45 journals, preprint servers and other sources, with the group of each source written next to its bar. Shades of grey identify the groups (darkest = largest group in A). The papers were discussed in sessions between June 2023 and June 2026. Journals were identified from the DOI prefix. One paper was discussed twice. Historical publications are classic papers (for example from 1920 and earlier).

The results of the sensitivity analyses are reported in the Supplementary Results (Supplementary Figures S1–S4, Tables S1–S5). The main themes recurred under alternative corpus definitions and clustering settings, although the number and boundaries of clusters were sensitive to the settings and to the random seed.

## Discussion

This study demonstrates how an archive created for open neuroscience education can also support quantitative investigation of scientific discourse. Across 470 transcripts spanning sixteen years, the CNSeminars archive captured changes in thematic attention, conceptual emphasis, methodological vocabulary, and discussion of open science. Short-term forecasting of vocabulary, by contrast, showed limited predictability: models improved on a persistence baseline mainly through smoothing, and recent growth in a term did not predict continued growth. These findings establish an additional analytical value for open seminar archives: alongside disseminating knowledge, they preserve a measurable record of how a scientific community frames its questions and directs its attention over time.

### Scientific discourse as a record of changing research priorities

Longitudinal text analysis offers established approaches for investigating changes in topics within document collections (Blei & Lafferty, 2006). The present study extends this perspective to an archive of spoken neuroscience. Whereas publications primarily document formal research outputs, seminar presentations also preserve how researchers explain their motivations, position findings within competing frameworks, and connect methodological developments to scientific questions. Analysing these exchanges therefore adds a complementary perspective on scientific change: the evolving language through which a community interprets its work.

The thematic organisation of CNSeminars reflects its origins in clinical neuroanatomy while documenting its expansion into a broader scientific forum (Figure 3). Language, white matter anatomy, connectivity, and clinical neuroscience remain prominent, alongside molecular mechanisms, computational approaches, and research practices. The enduring clinical-anatomical interests include questions about how language depends on distributed cortical, subcortical, and white matter systems (Forkel & Hagoort, 2024), and how connectional anatomy informs explanations of language impairment (Forkel et al., 2020). This combination is informative because scientific communities develop through interactions between established interests and newly introduced questions. The resulting thematic structure captures both the community’s enduring scientific interests and the breadth of questions introduced through its educational programming. Its longitudinal interpretation must therefore account for the decisions through which speakers, courses, and papers enter the archive.

This study develops an analytical use envisaged by the OHBM Time Machine initiative: openly available recordings can provide a historical resource for tracing the development of scientific concepts and methods (Wearn et al., 2025). It also extends the earlier qualitative review of Neuroccino discussions by examining temporal patterns across a substantially larger archive (Guzmán Chacón et al., 2024). The contribution is, therefore, substantive and methodological: the results characterise this community’s scientific conversations, while the analytical approach provides a basis for comparisons with other seminar archives.

### Changing conceptual emphasis within neuroscience

The increasing prominence of network-related vocabulary is consistent with the development of network neuroscience, which examines brain organisation through relationships between interacting elements across spatial and temporal scales (Bassett & Sporns, 2017). Within CNSeminars, network terminology became more frequent than localisation-related terminology (Fig. 6). This pattern suggests an increasing emphasis on connectivity and distributed organisation in the language used to explain brain function (Thiebaut De Schotten & Forkel, 2022). Related debates within Neuroccino questioned the explanatory limits of localisation and the privileged status of discrete brain areas (Hayden et al., 2026; Noble et al., 2024). Regional specialisation and network relationships nevertheless remain complementary levels of explanation, and the observed trajectories indicate a change in their relative prominence within discourse.

The increased emphasis on individual variability similarly places the archive within a scientific context in which differences between participants have become objects of investigation in their own right (Forkel et al., 2026). For example, functional connectome fingerprinting demonstrated that individual connectivity profiles can be sufficiently distinctive to identify participants across scanning sessions (Finn et al., 2015). Within the archive, the growing visibility of individual differences suggests increasing attention to how variation between people informs explanations of brain organisation, alongside continuing discussion of group-level patterns.

The computational trajectories require a related distinction between methodological adoption and conceptual change. Increased discussion of machine learning, prediction, or Bayesian approaches may reflect new analytical capabilities, changing research objectives, or interest in the assumptions underlying inference. These approaches also address different questions and should not be interpreted as a single successor to conventional statistics. The repeated crossovers between task-related and resting-state vocabulary further illustrate why scientific development is better characterised here as changing emphasis than as a sequence of replacements. Taken together, these patterns depict scientific development as the coexistence and recombination of explanatory approaches, with their relative prominence changing over time.

### Open science becomes part of scientific conversation

The increased frequency of open science vocabulary indicates that research practices themselves became more visible topics of scientific discussion. References to reproducibility, sharing, and statistical reform connect the archive to wider efforts to improve the transparency and reliability of neuroimaging research. These efforts have emphasised transparent methodological reporting, accessible data and code, and changes to research practices and incentives that support reproducibility (Poldrack et al., 2017).

The archive thus records changing attention to both scientific questions and the conditions under which evidence is produced and shared. Registered Reports exemplify this concern with research practices: study questions and methods undergo peer review before research is undertaken, with in-principle acceptance intended to reduce dependence of publication decisions on the results obtained (Chambers, 2013; Chambers & Tzavella, 2021). Vocabulary frequencies cannot establish that speakers preregistered studies, shared data, or reproduced published findings. They do, however, document the presence of these concerns within the community’s scientific conversation. The expansion of community and open science programming may have contributed to their increased prominence, illustrating how educational curation can create space for discussion of research norms alongside substantive findings.

### Limited predictability of vocabulary change

The forecasting analyses extend the study beyond retrospective description. Relative to a persistence baseline, the regression model reduced mean absolute error by approximately 12% for one-step forecasts and 17% for two-step forecasts across observed annual documents. This improvement was not specific to the model, however, a simple average of the last three observations performed slightly better, suggesting that the gain reflected smoothing of noisy annual values rather than successful prediction of their direction of change. Moreover, recent growth did not predict further growth. A score combining recent slope, acceleration, and recency of activity was negatively correlated with subsequent change at every forecast origin, meaning that the terms that had risen most tended to fall back. These results are consistent with mean reversion in vocabulary frequencies. Annual fluctuations may also reflect the speakers, courses, and papers selected for the channel, although the forecasting analyses do not establish the causes of those fluctuations.

Descriptive trajectories of vocabulary could nevertheless support educational planning. Sustained growth in methodological terminology might motivate tutorials, while increasing discussion of a conceptual question could inform a debate or focused seminar series. Such trajectories could therefore provide an additional input to curation, considered alongside scientific judgement and audience needs. Because short-term increases often reversed in the backtests, however, these trajectories describe recent attention rather than reliably forecasting future interests. Their practical value would require prospective evaluation, including whether they identify useful educational priorities and generalise across communities.

Interpretation of emerging-topic rankings also requires care. The ranking of terms by recent growth did not demonstrate predictive validity in temporal backtesting (Fig. 9), so it is reported as a description of recent interests rather than evidence of sustained topic emergence (Fig. 10). Established terms such as “neuron” or “fMRI” rank among the fastest growing terms without representing newly emerging concepts. Growth, recent acceleration, and conceptual novelty are distinct properties. Future analyses could distinguish the arrival of new terminology from renewed attention to established ideas and changes in the contexts in which those ideas are discussed. Comparisons with publication and conference-abstract corpora would further test whether seminar discourse precedes, follows, or develops alongside changes in formal scientific outputs.

### Educational reach, global inclusivity, and public responsibility

The channel’s lifetime viewing activity demonstrates that the archive has been accessed well beyond its original institutional setting. More than 203,000 views and approximately 17,000 hours of watch time document substantial use, while geographic reporting records viewing across continents, including countries of the Global South. Recent viewing across both live and recorded formats indicates that the channel supports different modes of participation. These findings demonstrate international reach and sustained use, providing a basis for evaluating how the archive serves audiences across research and educational settings.

Making expert instruction available without membership or subscription requirements removes one barrier to participation. Recordings also allow audiences to engage at different times and revisit material. Work on inclusive online neuroscience events emphasises that these opportunities must be accompanied by deliberate attention to accessibility, participation, and the needs of groups who may otherwise be excluded (Levitis et al., 2021). International availability alone does not establish equitable access, learning gains, or opportunities to contribute. For CNSeminars, advancing inclusion, therefore, involves attention to language, caption quality, connectivity, and representation among speakers, discussion leaders, and organisers.

The reciprocal exchange highlighted in the Introduction is particularly relevant to this next stage of development. An internationally accessible archive can broaden the circulation of expertise, while participatory formats can create opportunities for audiences to shape the questions and perspectives represented. Brainhack initiatives (e.g., https://youtube.com/playlist?list=PLg2e4R8Sdhpf2BxO5gB_Qkqo1Uv6pllJ-&si=7sgbOhYSEVoZeMLK, https://youtube.com/playlist?list=PLXaXn1qgv0q8&si=Bagvj3Z1zeClZDqU) and Brain Mappers of Tomorrow illustrate complementary approaches through collaborative learning and multilingual engagement, respectively (Gau et al., 2021; Lee et al., 2024). Future evaluation of CNSeminars could examine audience needs, barriers to participation, and whose expertise enters the programme, alongside geographic viewing patterns. Such evaluation would connect the channel’s commitment to openness with evidence about the inclusivity of its educational model.

The educational model also includes active contributions from students who prepare and lead Neuroccino journal club discussions. These activities create opportunities to practise critical appraisal, synthesis, presentation, and discussion with researchers across career stages and institutions. This participatory dimension complements the recorded archive: students contribute to the scientific conversation while developing skills in evaluating and communicating research. Assessing these experiences would provide a more direct measure of educational value than viewing statistics alone.

For publicly funded research, accessible communication is also a matter of societal responsibility. Sharing the methods, reasoning, and uncertainties of neuroscience makes expertise available to students, practitioners, adjacent disciplines, and interested members of the public. This responsibility extends beyond disseminating completed findings. It includes making the processes through which scientific knowledge is interpreted and debated more accessible.

### Recognising communication as scholarly infrastructure

The channel’s history illustrates what sustained community effort can produce, but continued provision requires time, expertise, and organisational support. Preparing discussions, supporting speakers, recording sessions, maintaining metadata, checking transcripts, and preserving materials are substantive scholarly activities. The OHBM Time Machine paper similarly documents the technical workflows, manual curation, and coordination involved in making recorded scientific content openly accessible (Wearn et al., 2025). The archive’s dual value as an educational resource and a corpus for research strengthens the case for recognising this work within academic workloads.

We, therefore, argue that science communication and educational curation should receive explicit recognition in workload allocation, training, and academic assessment. Dedicated teaching components, such as science communication training within postgraduate programmes, provide one route to developing these skills as part of the academic curriculum. Recognition should also extend to the students and researchers who sustain community resources, through appropriate credit, training, and allocated time. Where communication is expected in funded research, institutions should provide the support needed to deliver and maintain it. Embedding these responsibilities within research and teaching roles would improve continuity and reduce dependence on discretionary effort.

Preservation and reuse require infrastructure beyond public video hosting. Reliance on a commercial platform leaves continued access dependent on services and policies outside the scientific community’s control (Wearn et al., 2025). Channel removal or service discontinuation could make an archive inaccessible; without independent backups, its contents could be lost. The FAIR principles provide a framework for making digital research objects findable, accessible, interoperable, and reusable (Wilkinson et al., 2016). For seminar archives, these principles should be accompanied by preservation arrangements covering original recordings, versioned transcripts, metadata, permissions, and analysis code. YouTube can provide the public-facing interface, while an independently maintained repository safeguards the materials and analytical record. The scientific memory created through open education should outlast the platform through which it is shared.

### Limitations

The archive is curated, with coverage shaped by the organisers’ interests, speaker participation, and the selection of papers for Neuroccino. The papers discussed at Neuroccino sessions were drawn mainly from high-profile journals, with Nature Portfolio Group accounting for 35% of the selected papers (Fig. 10), so this component reflects both editorial selection within those outlets and the community’s reading choices. This selectivity is also what makes the archive informative: it records what one international neuroimaging community chose to discuss over time, which is the objective of study here, and makes no claim to represent the neuroscience literature as a whole. Temporal interpretation is constrained by unequal annual sample sizes (fewer than ten transcripts per year before 2020), missing recordings in 2018–2019, incomplete coverage of 2026, and a programme that changed substantially with the expansion of online activities around the COVID-19 pandemic. Trends before 2020 are therefore interpreted cautiously. Changes in scientific attention cannot be fully separated from changes in programming, so the trajectories are best read as shifts in what this community discussed.

Several features of the design bound the way the results should be interpreted. Automated transcription introduces uncertainty for specialist terminology, acronyms, and recordings of variable quality, but the analyses rest on aggregate patterns across 470 transcripts rather than on individual words or videos, which limits the influence of isolated transcription errors. Vocabulary frequencies measure explicit wording, and the keyword groups used for conceptual, methodological and open science analyses are proxies, so the trajectories describe associated vocabulary rather than the use of a particular method or paradigm. The forecasting analysis is a proof of concept on a short, unevenly sampled series, and its generalization to other achives requires further validation. Together, these data support an account of the channel’s international reach, while questions of educational effectiveness and equitable participation would need a dedicated study. Taken as a whole, these limitations define the scope of the conclusions rather than undermining them; the study offers a transparent and reproducible account of how one neuroimaging community’s discourse evolved.

### Towards a distributed observatory of scientific discourse

A next step would be to connect independently curated seminar archives through shared metadata and analytical standards. A distributed observatory of scientific discourse could compare communities, languages, and presentation formats while preserving the distinctiveness of each collection. Linking these archives to publications and conference abstracts would enable investigation of how ideas move between scientific conversations and formal outputs. Such comparisons would require harmonised transcript processing, explicit documentation of curation practices, and validation across languages and content types. Shared curation and preservation workflows would also allow grassroots initiatives to learn from one another and reduce duplicated effort. The present study provides an initial demonstration of the analytical opportunity: scientific exchanges preserved for education can become evidence for investigating how research communities develop their ideas. Sustaining and connecting these resources would support both their immediate educational mission and a broader understanding of scientific change.

## Conclusion

An open educational archive designed to disseminate neuroscience has become a research resource for examining how scientific communities organise and revise their ideas. Across sixteen years of CNSeminars, analyses of spoken discourse revealed changing thematic priorities, increasing emphasis on networks, individual variability, and computational approaches, and greater visibility of discussion about open science practices. Temporal backtesting further showed that smoothing improved prediction of annual vocabulary frequencies, while terms with the strongest recent growth tended to decline afterwards. These results limit the interpretation of recent vocabulary growth as evidence of sustained changes in scientific attention. Together, these findings establish a complementary approach to studying scientific change through the conversations in which researchers explain, contextualise, and debate their work.

Open neuroscience education creates a lasting record of the questions, explanations, and research practices that occupy a community’s attention. Recognising this record as scholarly infrastructure, with sustained support for inclusive participation, curation, and preservation, would strengthen its educational value and enable comparative investigation across communities. The conversations preserved today can become evidence for understanding how science develops over time.

## Code and data availability

All the transcripted videos and code used for the analyses are openly available at https://github.com/irenebellin/clinicalneuroanatomyseminars_10yearsanniversary.

## Acknowledgments

We thank Kurt Schilling for insightful discussions, and the extended CNS team for their contributions throughout the channel’s history. In particular, we acknowledge Sanja Budisavljevic, Michel Thiebaut de Schotten, Henrietta Howells, Lilit Dulyan, Muge Sertel Ozker, Edoardo Gornetti, Valentina Pacella, Hervé Lemaitre, Lauren Thiebaut, Marcela Ovando-Tallez, Eva Guzmán Chacón, Chris Foulon, Ansley Martin, Sjoerd Murris, Sabrina Beber, Anna Matsulevits, Lia Talozzi, Victor Nozais, Laura Vavassori, Patrick Friedrich, Sahba Besharati, Marco Catani, and Fergus Kane. A very special thank goes to Heidemarie Kranich, Carola Forkel, for their continuous support, feedback, and encouragement. We also gratefully acknowledge the support of NatBrainLab, King’s College London, the Wellcome Trust, and the BCBlab.

The archive exists because speakers have generously shared their expertise and students around the world have volunteered their time, curiosity, and energy to prepare Neuroccino discussions and participate in the sessions. We extend a special thank you to Luna Productions, Gary Weimberg and Catherine Ryan, for bringing Marian Diamond’s life and work to our community through what became the channel’s most watched content. Her commitment to sharing knowledge, captured in “Each one, teach one”, continues to inspire the channel’s educational mission. Lastly, we thank the members of the Clinical Neuroanatomy of Language research team at the Donders Institute and the Max Planck Institute for Psycholinguistics for their valuable feedback and sustained support.

## Author Contribution

I.B. conducted the primary analyses, code development, methodology, and generated the visualisations and interpretation of the findings. L.K.T. curated the integrated dataset and the full transcription of the dataset. A.B. set up the original YouTube channel in 2016 and has curated content ever since. A.W. contributed to the methodology, code development, analyses and interpretation. S.J.F. conceived the study, secured funding and provided overall supervision and project administration. All authors contributed to writing the original manuscript and to its subsequent review and editing, and approved the final version.

## Supplementary

### Supplementary Methods

#### Sensitivity analysis

The composition of the corpus changed markedly over time: 41 videos before 2018, none in 2018–2019, and 425 from 2020 onwards, with 239 of the 466 analysed videos being Neuroccino sessions. The 466 videos are the transcripts retained by the main clustering analysis (at least 50 tokens after preprocessing). We therefore tested whether the clustering in the main analysis depends on how the corpus is defined, sampled or analysed. In each scenario the full pipeline (TF-IDF vectorisation, UMAP, HDBSCAN) was re-run on the relevant subset with the main-text settings: 3,000 terms; cosine UMAP with 20 neighbours, a minimum distance of 0.05 and two dimensions; HDBSCAN with a minimum cluster size of 15 and a minimum of three samples.

The scenarios were: (i) all 466 analysed videos (baseline); (ii) conventional CNSeminars only (n = 85); (iii) excluding only Neuroccino (n = 227); (iv) excluding short-form content, defined as a raw word count below 1,500 (n = 441 retained) or below 3,000 words (n = 401 retained); (v) the dense period 2020–2026 only (n = 425); and (vi) year-balanced subsamples with at most 20 videos per year (n = 181), repeated in 30 random draws. For vocabulary-level results we also pooled the TF-IDF vectors with each video weighted by the inverse of the number of videos in its year.

#### Random seeds and seed-to-seed reference

UMAP is stochastic, so partitions differ between runs even on identical data. Every scenario was therefore repeated with ten random seeds and compared with the baseline run of the same seed. To quantify how much agreement seed variation alone allows, we compared baseline runs from neighbouring seeds (1 vs 2, 2 vs 3, …, 10 vs 1). The main analysis used seed 42. A solution with four or fewer clusters was called collapsed.

#### Minimum cluster size in subset

A fixed minimum cluster size of 15 is large relative to small subsets. Each subset scenario was therefore run twice: with the fixed value, and with the value scaled in proportion to the number of videos (15 × n / 466, rounded, minimum 5).

#### Agreement measures

Agreement between two partitions was measured with the adjusted Rand index (ARI; Hubert & Arabie, 1985) on the videos assigned to a cluster in both runs, reported as median and interquartile range (IQR) across seeds. Clusters were matched across runs by the overlap of their member videos (Jaccard index) using optimal one-to-one assignment (Kuhn, 1955). A baseline cluster was counted as reproduced when its Jaccard index was at least 0.5, and as partly reproduced between 0.3 and 0.5. For matched clusters with a Jaccard index of at least 0.3 we compared yearly prevalence trajectories (the share of each year’s videos falling in the cluster) with Spearman correlation, using years with at least three videos in the comparison run and requiring at least five such years.

#### Clustering parameters and dimensionality

With the main-text seed (42), we varied the minimum cluster size (8, 10, 15, 20, 30), minimum samples (1, 3, 5), UMAP neighbours (10, 15, 20, 30) and minimum distance (0, 0.05, 0.1) in a full grid of 180 settings, and the TF-IDF vocabulary size (1,500 and 5,000 terms) in two further runs. Because HDBSCAN can also be run on more than two UMAP dimensions, with the two-dimensional projection kept for visualisation, we additionally clustered in 2, 5 and 10 dimensions, crossed with neighbours (10, 15, 20, 30) and minimum cluster size (8, 10, 15, 20), using three seeds per configuration. Stability of a configuration was the mean pairwise ARI between its three seeds.

#### Cross-check with ten dimensional clustering

We clustered the videos in 10 UMAP dimensions with 30 neighbours and a minimum cluster size of 10 or 15 (ten seeds each), and compared the result with the clusters of the main analysis. Clusters of the seed that agreed most with the other seeds were matched to the main-analysis clusters as described above, and described by the terms whose TF-IDF weight in the cluster most exceeded the corpus mean. Cluster labels from this check were drawn on the unchanged two-dimensional map.

### Supplementary results

At seed 42 the baseline gave 9 clusters, with 116 of 466 videos (24.9%) not assigned to a cluster. Across seeds 1–10, the same settings gave between 2 and 14 clusters (median 7.5); four seeds gave a coarse solution (two gave two clusters, one gave three and one gave four). Agreement between baseline runs of neighbouring seeds was low (median ARI 0.10, IQR 0.04–0.23), whereas yearly prevalence trajectories of matched clusters were more stable (median Spearman ρ = 0.96). These values are the reference for all other comparisons: a scenario is considered to change the partition only if agreement falls clearly below them.

Agreement in the dense period (median ARI 0.34 with a fixed and 0.25 with a scaled minimum cluster size; Table S1, Fig S1) and with the 3,000-word short-form cut-off (0.31 and 0.49) was higher than agreement between baseline runs of neighbouring seeds (0.10), as was the 1,500-word-cut-off (0.26 and 0.31), although the latter was more variable (IQR 0.05-0.49). Excluding Neuroccino (227 videos) gave a median ARI of 0.36 (fixed) and 0.13 (scaled). Keeping only the 85 CNSeminars removes most of the corpus and with it whole clusters, so low agreement with the full partition is expected (median ARI 0.03 and 0.09); they are too few to cluster at this resolution (median 2 clusters with a fixed and 4.5 with a scaled minimum cluster size). With the fixed minimum cluster size, 100% of runs collapsed to four or fewer clusters for CNSeminars only and for the corpus without Neuroccino, 87% for year-balanced draws and 60% below 1,500 words; scaling the minimum cluster size reduced these to 50%, 60%, 30% and 40%. Yearly prevalence trajectories of matched clusters stayed correlated (median ρ 0.74–0.88) in the dense-period, short-form and Neuroccino-excluded scenarios, but not for CNSeminars only (−0.02 and 0.39) or year-balanced draws (0.54-0.56).

**Table S1.** Agreement with the baseline partition (same seed) over seeds 1–10. Each cell shows fixed / scaled minimum cluster size; the first row is the seed-to-seed (floor) reference.

| Scenario | Videos | Clusters, median | Collapsed runs | ARI, median (IQR) | Trajectory $\rho$, median |
| --- | --- | --- | --- | --- | --- |
| Baseline vs neighbouring seed (floor) | 466 | 7.5 | 40% | 0.10 (0.04–0.23) | 0.96 |
| CNSeminars only | 85 | 2 / 4.5 | 100% / 50% | 0.03 (0.00–0.14) / 0.09 (0.00–0.27) | -0.02 / 0.39 |
| Excluding Neuroccino | 227 | 2 / 3 | 100% / 60% | 0.36 (0.09–0.72) / 0.13 (0.09–0.53) | 0.87 / 0.85 |
| Excluding short-form (< 1,500 words) | 441 | 2.5 / 5 | 60% / 40% | 0.26 (0.06–0.42) / 0.31 (0.05–0.49) | 0.82 / 0.81 |
| Excluding short-form (< 3,000 words) | 401 | 7.5 / 9 | 20% / 20% | 0.31 (0.16–0.59) / 0.49 (0.19–0.61) | 0.75 / 0.74 |
| 2020–2026 only | 425 | 8.5 / 8.5 | 30% / 30% | 0.34 (0.07–0.62) / 0.25 (0.11–0.60) | 0.86 / 0.88 |
| Year-balanced, cap 20 | 181 | 3 / 9 | 87% / 30% | 0.16 (0.05–0.43) / 0.11 (0.04–0.38) | 0.56 / 0.54 |

**Figure S1.**
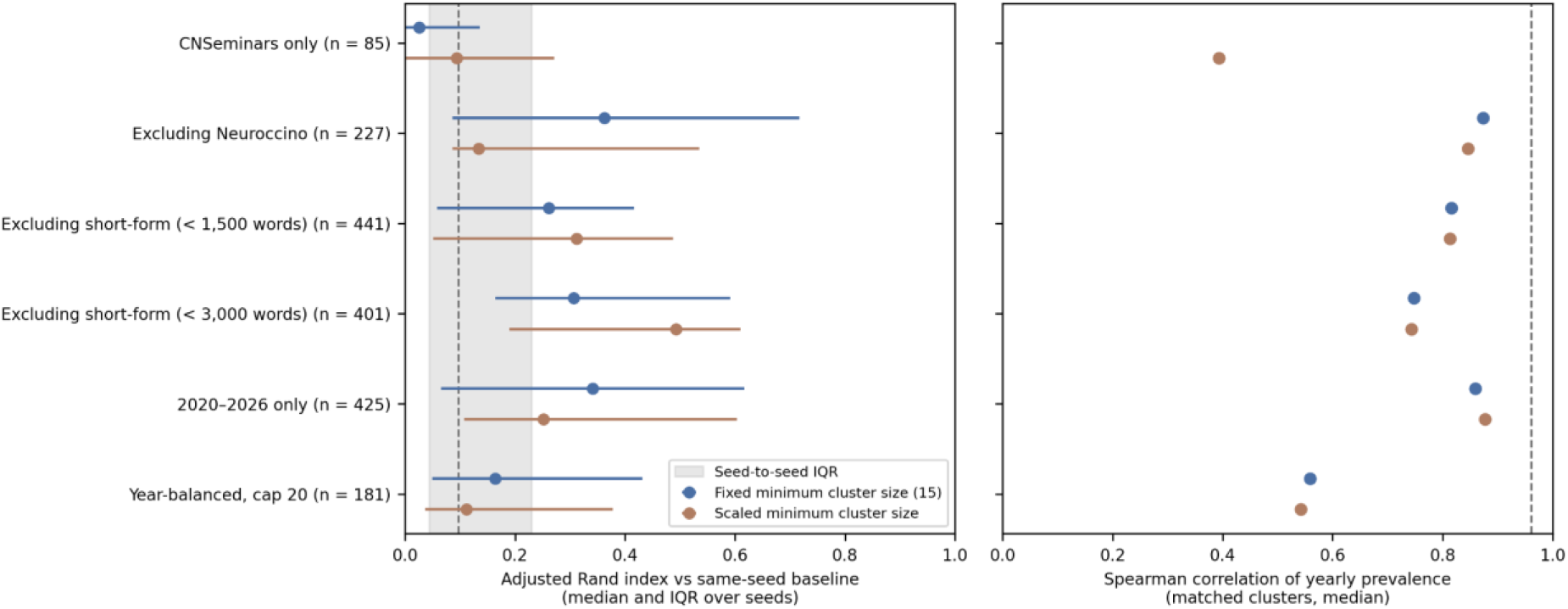
Agreement with the baseline partition under alternative corpus definitions. Left: adjusted Rand index (median and interquartile range over ten seeds; 30 draws for the year-balanced scenario) against the baseline run of the same seed, for a fixed and a scaled minimum cluster size; the grey band is the interquartile range of agreement between baseline runs of neighbouring seeds and the dashed line its median. Right: median Spearman correlation of yearly prevalence of matched clusters; the dashed line is the seed-to-seed median.

In year-balanced draws with the fixed minimum cluster size, 87% of runs collapsed to four or fewer clusters. This reflects the 181-video sample rather than its content: scaling the minimum cluster size gave a median of 9 clusters (coarse solutions 30%, median ARI 0.11, trajectory ρ 0.54; Table S1). Weighting each video by the inverse of its year’s video count changed the pooled vocabulary moderately: the ranking of all 3,000 terms was preserved (Spearman ρ = 0.80), but 62 of the top 100 terms were shared (Jaccard index 0.45). Pooled term rankings therefore describe the corpus as uploaded, which is dominated by 2020–2026.

In the 180-setting grid (seed 42; Table S2, Fig S2), the number of clusters depended on the number of UMAP neighbours and on the minimum cluster size, with a transition between fine and coarse solutions. Across the grid, 38% of settings (69 of 180) gave four or fewer clusters. With 10 or 15 neighbours the median solution had 7 or 3 clusters (median ARI 0.16 and 0.06 against the baseline); with 20 neighbours it had 9 clusters (median ARI 0.51), and with 30 neighbours 9 clusters (0.55). Median cluster numbers fell from 21 at a minimum cluster size of 8 to 15 at 10, 7 at 15, 3 at 20 and 2 at 30. Changing the TF-IDF vocabulary to 1,500 terms gave 9 clusters (ARI 0.59), whereas 5,000 terms gave a collapsed two-cluster solution (ARI 0.06; one run each), so the baseline partition seems sensitive to the vocabulary size.

**Table S2.** Parameter grid summarised by parameter value (seed 42). For each value, the other parameters vary across the grid. ARI is against the main-analysis partition; a solution with four or fewer clusters counts as coarse. The TF-IDF rows are single runs at the main-text settings.

| Parameter | Value | Settings | Clusters, median | Clusters, range | ARI vs baseline, median | Collapsed settings |
| --- | --- | --- | --- | --- | --- | --- |
| UMAP neighbours | 10 | 45 | 7 | 2–28 | 0.16 | 33% |
| UMAP neighbours | 15 | 45 | 3 | 2–29 | 0.06 | 69% |
| UMAP neighbours | 20 | 45 | 9 | 2–26 | 0.51 | 27% |
| UMAP neighbours | 30 | 45 | 9 | 2–28 | 0.55 | 24% |
| Minimum cluster size | 8 | 36 | 21 | 2–29 | 0.39 | 3% |
| Minimum cluster size | 10 | 36 | 15 | 2–21 | 0.45 | 14% |
| Minimum cluster size | 15 | 36 | 7 | 2–13 | 0.35 | 39% |
| Minimum cluster size | 20 | 36 | 3 | 2–10 | 0.15 | 58% |
| Minimum cluster size | 30 | 36 | 2 | 2–7 | 0.06 | 78% |
| Minimum samples | 1 | 60 | 10.5 | 2–29 | 0.38 | 25% |
| Minimum samples | 3 | 60 | 8 | 2–28 | 0.34 | 43% |
| Minimum samples | 5 | 60 | 5.5 | 2–22 | 0.24 | 47% |
| Minimum distance | 0 | 60 | 10 | 2–29 | 0.38 | 37% |
| Minimum distance | 0.05 | 60 | 7 | 2–28 | 0.32 | 35% |
| Minimum distance | 0.1 | 60 | 6.5 | 2–26 | 0.35 | 43% |
| TF-IDF vocabulary (single run) | 1,500 | 1 | 9 | 9–9 | 0.59 | 0% |
| TF-IDF vocabulary (single run) | 3,000 | 1 | 9 | 9–9 | 1.00 | 0% |
| TF-IDF vocabulary (single run) | 5,000 | 1 | 2 | 2–2 | 0.06 | 100% |

Clustering in more dimensions was more stable (Fig S2, bottom; Table S3). Mean between-seed ARI within a configuration had a median of 0.49 for two dimensions (range 0.15–0.75), 0.75 for five (0.31–0.91) and 0.84 for ten (0.61–0.94). At least one of the three seeds collapsed in 56% of two-dimensional, 31% of five-dimensional and 6% of ten-dimensional configurations. The main-text setting (20 neighbours, minimum cluster size 15) was unstable in two dimensions (between-seed ARI 0.15) and five-dimensions (0.32; one of three seeds collapsed in both) but stable in ten (0.81, no collapsed seed), whereas 30 neighbours was more stable throughout (median 0.58 in two, 0.75 in five and 0.85 in ten dimensions).

The most stable configuration used ten dimensions, 10 neighbours and a minimum cluster size of 20 (ARI 0.94, 8-10 clusters, no collapsed seeds).

**Figure S2.**
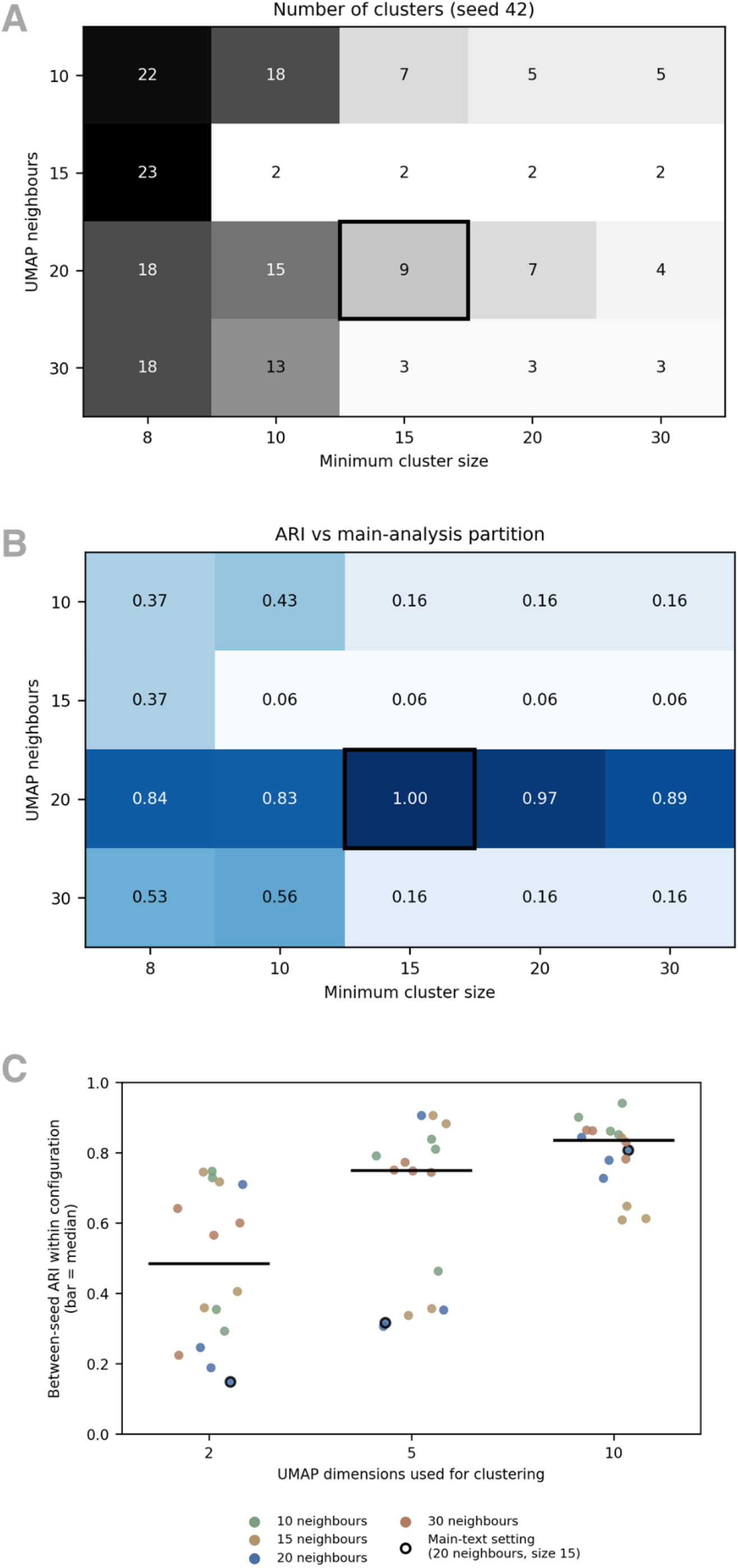
Sensitivity to clustering parameters. Top and middle: number of clusters and ARI against the main-analysis partition by UMAP neighbours and minimum cluster size (minimum samples 3, minimum distance 0.05, seed 42); the box marks the main-text setting. Bottom: mean between-seed ARI within each configuration (three seeds) by the number of UMAP dimensions used for clustering; bars mark medians and black-edged points the main-text setting.

**Table S3.** Stability of clustering by the number of UMAP dimensions used. Between-seed ARI is the mean pairwise ARI between three seeds within a configuration (neighbours × minimum cluster size); the last four columns give the median over minimum cluster sizes for each number of neighbours.

| UMAP dimensions | Configurations | Between-seed ARI, median | Between-seed ARI, range | Configurations with a collapsed seed | ARI, 10 neighbours | ARI, 15 neighbours | ARI, 20 neighbours | ARI, 30 neighbours |
| --- | --- | --- | --- | --- | --- | --- | --- | --- |
| 2 | 16 | 0.49 | 0.15-0.75 | 56% | 0.54 | 0.56 | 0.22 | 0.58 |
| 5 | 16 | 0.75 | 0.31-0.91 | 31% | 0.8 | 0.62 | 0.33 | 0.75 |
| 10 | 16 | 0.84 | 0.61-0.94 | 6% | 0.88 | 0.63 | 0.79 | 0.85 |

Clustering in ten dimensions with 30 neighbours gave no coarse solutions: 9-16clusters at a minimum cluster size of 10 and 8–10 at 15. Agreement between seeds was high (mean ARI 0.85 and 0.86). Agreement with the main-analysis partition was moderate (median ARI 0.55 and 0.52) and lower than that between the ten-dimensional seeds, so the main analysis partition and the more stable ten-dimensional solutions share their main themes but differ in their boundaries. Matched clusters had a median trajectory correlation of 0.84 and 0.78. Both solutions are drawn on the unchanged two-dimensional map in Fig S4.

At a minimum cluster size of 10 (Table S4, Fig S3), three of the nine main-analysis clusters were recovered one-to-one (Jaccard index ≥ 0.5): genetics and disease (T1), open science and reproducibility (T2), and language and speech (T8). Four were partly recovered (Jaccard 0.32–0.41): evolution and comparative anatomy (T3), diffusion MRI and axons (T5), networks, lesions and connectivity (T7), and language, lesions and white matter (T9). T7 appears to have split into a network and connectivity cluster (45 videos) and a lesion and disconnection cluster (31 videos). Two clusters were not recovered as single clusters: cerebellum and gross anatomy (T4) and the largest cluster, cellular and molecular neuroscience (T6, 89 videos), whose best match was a 16-video cluster on neurons and receptors; a new cluster of 12 videos on cells and glia appeared. Three new clusters of 12, 31 and 13 videos appeared in total (cells and glia; lesion, disconnection and stroke; and social, career and mentoring themes). At a minimum cluster size of 15 the solution was coarser (Table S5): two clusters recovered (T1, T2), three partly (T5, T7, T8) and four not recovered (T3, T4, T6, T9).

**Table S4.**
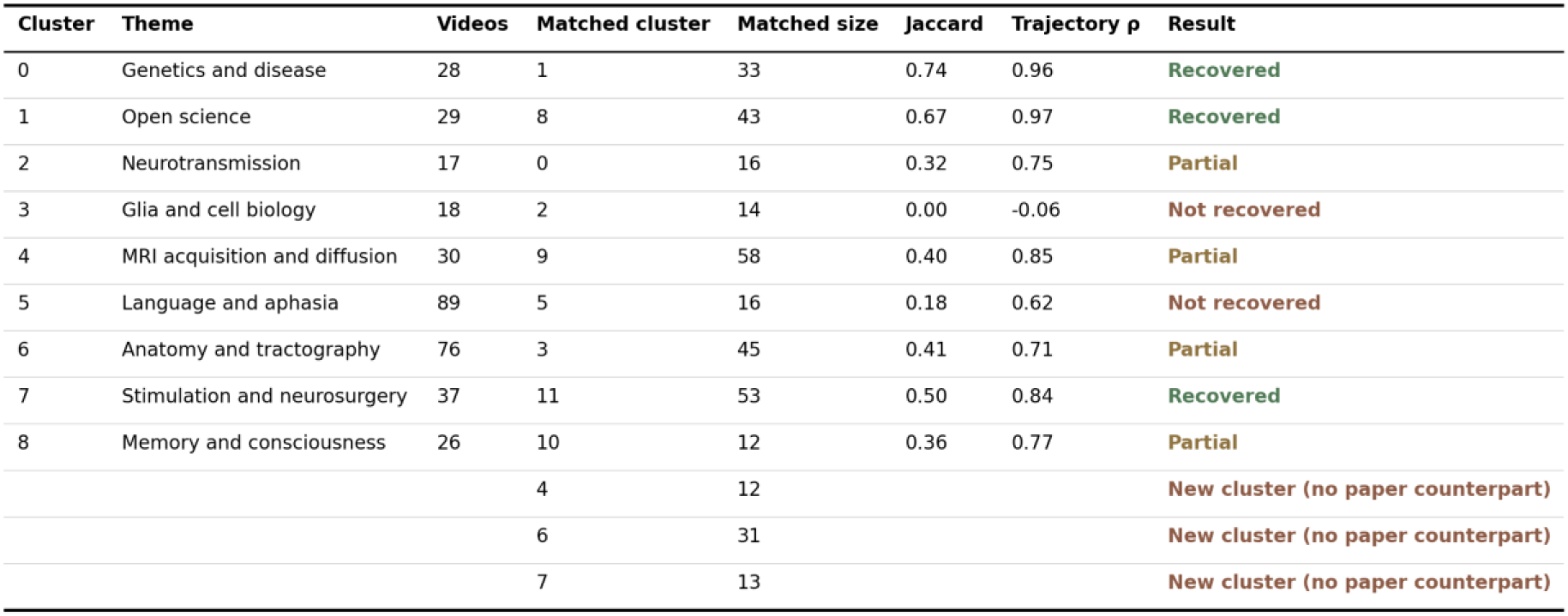
Main-analysis clusters (T1-T9 in the main text) matched to the ten-dimensional solution (30 neighbours, minimum cluster size 10, most representative seed). Theme names are descriptive labels from top terms.

**Figure S3.**
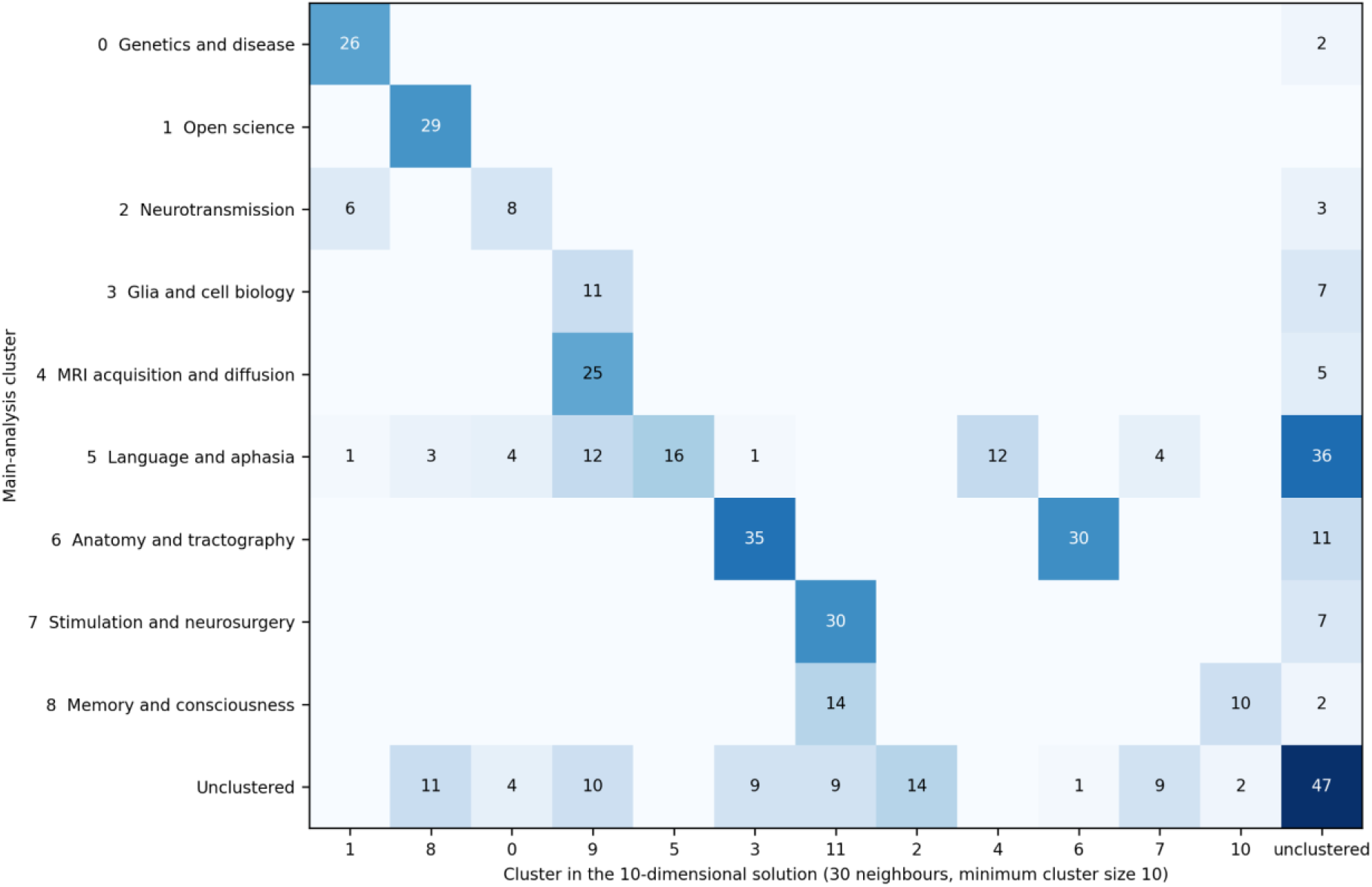
Overlap between main-analysis clusters (rows) and the ten-dimensional solution (columns; 30 neighbours, minimum cluster size 10, most representative seed). Cells give numbers of videos; columns are ordered to follow the best match of each row.

**Table S5.** Same as Table S4, at a minimum cluster size of 15.

| Cluster | Theme | Videos | Matched cluster | Matched size | Jaccard | Trajectory $\rho$ | Result |
| --- | --- | --- | --- | --- | --- | --- | --- |
| 0 | Genetics and disease | 28 | 2 | 35 | 0.75 | 0.97 | Recovered |
| 1 | Open science | 29 | 0 | 42 | 0.69 | 0.95 | Recovered |
| 2 | Neurotransmission | 17 | 1 | 19 | 0.00 | 0.37 | Not recovered |
| 3 | Glia and cell biology | 18 | 4 | 67 | 0.00 | -0.05 | Not recovered |
| 4 | MRI acquisition and diffusion | 30 | 7 | 69 | 0.36 | 0.82 | Partial |
| 5 | Language and aphasia | 89 | 3 | 17 | 0.19 | 0.62 | Not recovered |
| 6 | Anatomy and tractography | 76 | 5 | 32 | 0.38 | 0.54 | Partial |
| 7 | Stimulation and neurosurgery | 37 | 6 | 90 | 0.38 | -0.01 | Partial |
| 8 | Memory and consciousness | 26 |  |  |  |  | Not recovered |

**Figure S4.**
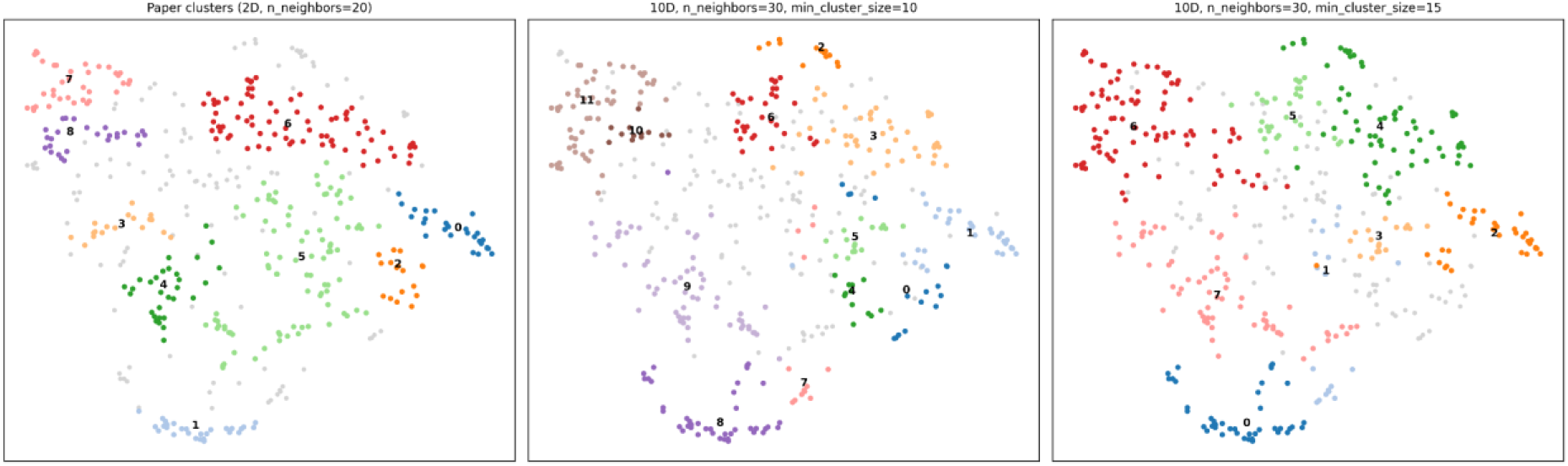
Two-dimensional UMAP map of the videos coloured by the main-analysis clusters (left) and by the ten-dimensional solutions at minimum cluster size 10 (middle) and 15 (right); grey points are unclustered.

### Summary of sensitivity analyses

Two findings stand out. First, restricting the corpus to the dense period or removing the shortest transcripts changed the partition no more than a different random seed did (median ARI 0.25–0.49 against 0.10 between neighbouring seeds), and the yearly trends of matched clusters persisted (median ρ 0.74–0.88). Second, the number and boundaries of clusters depend on the clustering settings and on the random seed: the baseline solution itself ranged from 2 to 14 clusters across seeds, the number of clusters changes with the number of UMAP neighbours and the minimum cluster size, and agreement between runs is higher when clustering in more dimensions. A ten-dimensional clustering recovered three of the nine themes closely (genetics and disease, open science, and language and speech) and four more partly, whereas the cerebellum and gross-anatomy cluster and the large cellular cluster were not recovered as single clusters. We therefore treat the main-analysis clusters as one partition of a continuous thematic landscape and base our conclusions on themes and their trends rather than on individual cluster boundaries or counts.

## References

Bassett, D. S., & Sporns, O. (2017). Network neuroscience. Nature Neuroscience, 20(3), 353–364. 10.1038/nn.4502

Bellier, L., Llorens, A., Marciano, D., Gunduz, A., Schalk, G., Brunner, P., & Knight, R. T. (2023). Music can be reconstructed from human auditory cortex activity using nonlinear decoding models. PLOS Biology, 21(8), e3002176. 10.1371/journal.pbio.3002176

Besharati, S., & Akinyemi, R. (2023). Accelerating African neuroscience to provide an equitable framework using perspectives from West and Southern Africa. Nature Communications, 14(1), 8107. 10.1038/s41467-023-43943-3

Blei, D. M., & Lafferty, J. D. (2006). Dynamic topic models. Proceedings of the 23rd International Conference on Machine Learning - ICML ’06, 113–120. 10.1145/1143844.1143859

Boulant, N., Mauconduit, F., Gras, V., Amadon, A., Le Ster, C., Luong, M., Massire, A., Pallier, C., Sabatier, L., Bottlaender, M., Vignaud, A., & Le Bihan, D. (2024). In vivo imaging of the human brain with the Iseult 11.7-T MRI scanner. Nature Methods, 21(11), 2013–2016. 10.1038/s41592-024-02472-7

Breslyn, W., & Green, A. E. (2022). Learning science with YouTube videos and the impacts of Covid-19. Disciplinary and Interdisciplinary Science Education Research, 4(1), 13. 10.1186/s43031-022-00051-4

Brossard, D. (2013). New media landscapes and the science information consumer. Proceedings of the National Academy of Sciences, 110(supplement_3), 14096–14101. 10.1073/pnas.1212744110

Campello, R. J. G. B., Moulavi, D., & Sander, J. (2013). Density-Based Clustering Based on Hierarchical Density Estimates. In J. Pei, V. S. Tseng, L. Cao, H. Motoda, & G. Xu (Eds.), Advances in Knowledge Discovery and Data Mining (Vol. 7819, pp. 160–172). Springer Berlin Heidelberg. 10.1007/978-3-642-37456-2_14

Chambers, C. D. (2013). Registered Reports: A new publishing initiative at Cortex. Cortex, 49(3), 609–610. 10.1016/j.cortex.2012.12.016

Chambers, C. D., & Tzavella, L. (2021). The past, present and future of Registered Reports. Nature Human Behaviour, 6(1), 29–42. 10.1038/s41562-021-01193-7

Finn, E. S., Shen, X., Scheinost, D., Rosenberg, M. D., Huang, J., Chun, M. M., Papademetris, X., & Constable, R. T. (2015). Functional connectome fingerprinting: Identifying individuals using patterns of brain connectivity. Nature Neuroscience, 18(11), 1664–1671. 10.1038/nn.4135

Frances, C., Forkel, S.J., Martin, A. E., Rowland, C. F. (2026). A European perspective on supporting research talent in academic institutions. Accepted in Nature Neuroscience.

Forkel, S. J., Dulyan, L., Schilling, K., & Thiebaut De Schotten, M. (2026). Neurovariability as a signature of adaptive brain function. Brain Structure and Function, 231(6), 83. 10.1007/s00429-026-03138-0

Forkel, S. J., & Hagoort, P. (2024). Redefining language networks: Connectivity beyond localised regions. Brain Structure and Function, 229(9), 2073–2078. 10.1007/s00429-024-02859-4

Forkel, S. J., Rogalski, E., Drossinos Sancho, N., D’Anna, L., Luque Laguna, P., Sridhar, J., Dell’Acqua, F., Weintraub, S., Thompson, C., Mesulam, M.-M., & Catani, M. (2020). Anatomical evidence of an indirect pathway for word repetition. Neurology, 94(6). 10.1212/WNL.0000000000008746

Gau, R., Noble, S., Heuer, K., Bottenhorn, K. L., Bilgin, I. P., Yang, Y.-F., Huntenburg, J. M., Bayer, J. M. M., Bethlehem, R. A. I., Rhoads, S. A., Vogelbacher, C., Borghesani, V., Levitis, E., Wang, H.-T., Van Den Bossche, S., Kobeleva, X., Legarreta, J. H., Guay, S., Atay, S. M., … Zuo, X.-N. (2021). Brainhack: Developing a culture of open, inclusive, community-driven neuroscience. Neuron, 109(11), 1769–1775. 10.1016/j.neuron.2021.04.001

Guzmán Chacón, E., Ovando-Tellez, M., Thiebaut De Schotten, M., & Forkel, S. J. (2024). Embracing digital innovation in neuroscience: 2023 in review at NEUROCCINO. Brain Structure and Function, 229(2), 251–255. 10.1007/s00429-024-02768-6

Hayden, B. Y., Heilbronner, S. R., & Yoo, S. B. M. (2026). Rethinking the centrality of brain areas in understanding functional organization. Nature Neuroscience, 29(2), 267– 278. 10.1038/s41593-025-02166-z

Hutchins, J. B., & Harris, A. (2025). Open Educational Resources for Undergraduate Neuroscience Education. Journal of Undergraduate Neuroscience Education, 23(2), e22–e25. 10.59390/JDTX8529

Lee, K., Borghesani, V., De Moraes, F., Pozzobon, A., Olsen, R., Kam, J. W. Y., Tzovara, A., & Badhwar, A. (2024). Brain Mappers of Tomorrow: An international multilingual initiative for neuroscience dissemination. Aperture Neuro, 4. 10.52294/001c.123400

Levitis, E., van Praag, C. D. G., Gau, R., Heunis, S., DuPre, E., Kiar, G., Bottenhorn, K. L., Glatard, T., Nikolaidis, A., Whitaker, K. J., Mancini, M., Niso, G., Afyouni, S., Alonso-Ortiz, E., Appelhoff, S., Arnatkeviciute, A., Atay, S. M., Auer, T., Baracchini, G., … Maumet, C. (2021). Centering inclusivity in the design of online conferences—An OHBM–Open Science perspective. GigaScience, 10(8), giab051. 10.1093/gigascience/giab051

McInnes, L., Healy, J., Saul, N., & Großberger, L. (2018). UMAP: Uniform Manifold Approximation and Projection. Journal of Open Source Software, 3(29), 861. 10.21105/joss.00861

Noble, S., Curtiss, J., Pessoa, L., & Scheinost, D. (2024). The tip of the iceberg: A call to embrace anti-localizationism in human neuroscience research. Imaging Neuroscience, 2, imag–2–00138. 10.1162/imag_a_00138

Nozais, V., Forkel, S. J., Foulon, C., Petit, L., & Thiebaut De Schotten, M. (2021). Functionnectome as a framework to analyse the contribution of brain circuits to fMRI. Communications Biology, 4(1), 1035. 10.1038/s42003-021-02530-2

Pedregosa, F., Varoquaux, G., Gramfort, A., Michel, V., Thirion, B., Grisel, O., Blondel, M., Müller, A., Nothman, J., Louppe, G., Prettenhofer, P., Weiss, R., Dubourg, V., Vanderplas, J., Passos, A., Cournapeau, D., Brucher, M., Perrot, M., & Duchesnay, É. (2012). Scikit-learn: Machine Learning in Python. 10.48550/ARXIV.1201.0490

Poldrack, R. A., Baker, C. I., Durnez, J., Gorgolewski, K. J., Matthews, P. M., Munafò, M. R., Nichols, T. E., Poline, J.-B., Vul, E., & Yarkoni, T. (2017). Scanning the horizon: Towards transparent and reproducible neuroimaging research. Nature Reviews Neuroscience, 18(2), 115–126. 10.1038/nrn.2016.167

Radford, A., Kim, J. W., Xu, T., Brockman, G., McLeavey, C., & Sutskever, I. (2022). *Robust Speech Recognition via Large-Scale Weak Supervision* (Version 1). arXiv. 10.48550/ARXIV.2212.04356

Silva, A. B., Liu, J. R., Zhao, L., Levy, D. F., Scott, T. L., & Chang, E. F. (2022). A Neurosurgical Functional Dissection of the Middle Precentral Gyrus during Speech Production. The Journal of Neuroscience, 42(45), 8416–8426. 10.1523/JNEUROSCI.1614-22.2022

Thiebaut De Schotten, M., & Forkel, S. J. (2022). The emergent properties of the connected brain. Science, 378(6619), 505–510. 10.1126/science.abq2591

Tournier, J., Mori, S., & Leemans, A. (2011). Diffusion tensor imaging and beyond. Magnetic Resonance in Medicine, 65(6), 1532–1556. 10.1002/mrm.22924

Wearn, A., Sitek, K. R., Valk, S. L., & Forkel, S. J. (2025). The Organization for Human Brain Mapping Time Machine: A freely accessible archive of Annual Meeting talks on YouTube. Aperture Neuro, 5(SI 2). 10.52294/001c.138647

Wilkinson, M. D., Dumontier, M., Aalbersberg, Ij. J., Appleton, G., Axton, M., Baak, A., Blomberg, N., Boiten, J.-W., Da Silva Santos, L. B., Bourne, P. E., Bouwman, J., Brookes, A. J., Clark, T., Crosas, M., Dillo, I., Dumon, O., Edmunds, S., Evelo, C. T., Finkers, R., … Mons, B. (2016). The FAIR Guiding Principles for scientific data management and stewardship. Scientific Data, 3(1), 160018. 10.1038/sdata.2016.18

World Wide Neuro. (n.d.). World Wide Neuro. Retrieved October 3, 2026, from https://www.world-wide.org/Neuro/

